# Nanomechanical phenotyping uncovers a non-canonical smooth muscle-like hypercontractility program in lung squamous cell carcinoma myofibroblasts

**DOI:** 10.64898/2026.11.06.764840

**Authors:** Annalisa Calò, Alejandro Bernardo, Natalia Díaz-Valdivia, Anna Labernadie, Victoria Batto, José Rodriguez-Rojas, Miguel Vizoso, Beatriz Cantero, Marselina Arshakyan, Marc Rico-Pastó, Larissa Huetter, Amelia Parker, Mohit Kumar, Núria Gavara, Gabriel Gomila, Xavier Trepat, Miguel Vicente-Manzanares, Jordi Alcaraz

## Abstract

Myofibroblast-like cancer-associated fibroblasts (myCAFs) drive the stiff, tumor-promoting microenvironment, yet their mechanical phenotype in non-small cell lung cancer (NSCLC) remains undefined. myCAFs from major NSCLC subtypes (lung adenocarcinoma (LUAD) and squamous cell carcinoma (LUSC)) were examined combining atomic force microscopy and traction force microscopy with molecular and signaling analyses. LUAD-myCAFs displayed canonical myofibroblast features, including elevated stiffness, traction forces, ROCK-MYPT1 signaling, and α-SMA-rich stress fibers. Unexpectedly, LUSC-myCAFs amplified these nanomechanical features despite reduced ROCK-MYPT1 signaling and comparable α-SMA levels. Instead, they exhibited a smooth muscle cell (SMC)-like contractile program characterized by enhanced SM-MHC(*MYH11*) expression colocalized with α-SMA, ERK1/2-RLC hyperactivation, and DNA methylation reprogramming, including SMC-contractility regulators. LUSC-myCAF hypercontractility was linked with reduced proliferative capacity and smaller intratumoral vessel lumens, consistent with the low CAF density and enhanced hypoxia patterns reported in LUSC. SMC markers were enriched and associated with adverse outcomes selectively in LUSC, highlighting the relevance of SMC-like myCAFs in LUSC progression. These findings unveil subtype-specific myCAF programs in NSCLC, and identify α-SMA⁺/SM-MHC(*MYH11*)⁺ myCAFs as key determinants of tumor mechanics, providing a rationale for the survival benefits of antihypertensive drugs targeting SMC hypercontractility. Moreover, they demonstrate that nanomechanical phenotyping can uncover translationally relevant myCAF phenotypes invisible to molecular profiling.

## 1. Introduction

Lung cancer is the leading cause of cancer-related mortality worldwide, with tobacco smoking as its major risk factor ^[1,2]^. Most cases are diagnosed as non-small cell lung cancer (NSCLC), with lung adenocarcinoma (LUAD) and lung squamous cell carcinoma (LUSC) being the most prevalent histological subtypes ^[3]^. LUAD and LUSC also differ in anatomic, molecular and clinical characteristics, which influence disease evolution and therapeutic responses ^[1,4,5]^. Although both subtypes originate from epithelial cells, increasing evidence highlights the general contribution of the surrounding tumor stroma to cancer progression and therapeutic resistance ^[6,7]^. The tumor stroma is highly fibrotic/desmoplastic in NSCLC and other solid tumors, a condition driven by the accumulation of cancer-associated fibroblasts (CAFs), which promote the abundant deposition of fibrillar collagens ^[8,9]^. Most CAFs in NSCLC exhibit an activated/myofibroblast-like phenotype (referred to as myCAFs) ^[10–13]^ traditionally identified by the expression of the contractility cytoskeleton marker alpha-smooth muscle actin (α-SMA). Remarkably, high α-SMA expression has been associated with immunosuppression and poor prognosis in NSCLC ^[8,11]^, supporting a major role for myCAFs in tumor progression. Likewise, TGF-β1, the most potent inducer of the myofibroblast phenotype ^[14,15]^, is frequently overexpressed in NSCLC ^[16]^, where it correlates with adverse outcome and drives the pathologic accumulation of myCAFs ^[17]^, making it a suitable factor to model desmoplasia in culture ^[7]^.

A major functional consequence of desmoplasia is the chronic stiffening that is a hallmark of solid tumors ^[18,19]^, which is emerging as a major driver of tumor progression through the enhancement of cancer cell proliferation, invasion and dissemination ^[20]^, CAF density ^[10]^, and fibrosis ^[15,21]^. Elevated mechanical forces in tumors can also compress blood vessels, exacerbating hypoxia and hindering drug delivery ^[19]^, and may foster immunosuppression by opposing T-cell infiltration ^[11,18]^. These broad tumor-promoting consequences underscore the need to define the aberrant tumor mechanics and elucidate its underlying mechanisms, to ultimately help develop therapeutic strategies preventing and/or reversing their associated pathological effects ^[14,18]^. Notably, myCAFs have been pointed as major drivers of tumor stiffening ^[18,20]^, owing to the enhanced contractility associated with α-SMA positive (α-SMA^+^) fibroblasts in tissue repair and organ fibrosis ^[14]^, particularly when α-SMA is incorporated into stress fibers ^[15,22,23]^. However, CAF mechanobiology is likely to be complex, given the emerging heterogeneity of CAFs in solid tumors ^[11,24]^, which is strongly influenced by tumor histology in NSCLC ^[7,11,25]^. Yet, whether and how this histotype-dependent heterogeneity influences the mechanical properties of myCAFs remains unknown.

The mechanical properties of cells depend both on passive cell material properties and active force generation, a process largely dependent on the contractile actomyosin cytoskeleton ^[26,27]^. In non-muscle cells such as fibroblasts and myofibroblasts, contractility arises mainly from the mechanical interactions between the actin cytoskeleton and non-muscle myosin II (NMMII) motors, whose activity is largely regulated by Rho-associated kinase (ROCK)-signaling ^[15,28,29]^. Two nanotechniques have been extensively used to study cell mechanics at physiologically-relevant length scales (nanometer to micrometer): atomic force microscopy (AFM) and traction force microscopy (TFM) ^[27,30]^. These nanotechniques provide complementary information on the nanomechanical phenotype of cells at the intracellular (AFM) and extracellular (TFM) level, by probing both shared features (active actomyosin contractility) and technique-specific cellular structures (cell cortex and underlying cytoskeleton in AFM ^[31,32]^, extracellular force transmission through focal adhesions in TFM ^[26]^). However, previous cell nanomechanics studies in cancer have focused predominantly on cancer cells, using these techniques in isolation ^[30,33]^. In contrast, our current understanding of the mechanics of CAFs, and of myCAFs specifically, is very limited, particularly in NSCLC ^[34–36]^. Here, we aimed to determine the nanomechanical characteristics of myCAFs from major NSCLC histologic subtypes by combining AFM and TFM, to identify key underlying processes, and to elucidate potential implications for tumor progression.

## 2. Results

### 2.1. AFM and TFM consistently reveal a coordinated three-fold increase in stiffness and traction forces in control pulmonary fibroblasts stimulated with TGF-β1

Histologic and single-cell transcriptomic/proteomic analyses have revealed that most lung CAFs are α-SMA^+ [10–13]^, and that cultured CAFs express more α-SMA than control fibroblasts (CFs) from uninvolved pulmonary tissue ^[12,36,37]^. To reduce phenotypic heterogeneity and enable meaningful mechanical comparisons between CAFs and CFs, all fibroblasts were cultured at high seeding density approximating that of histologic sections ^[10,17]^ (**Figure S**1) and stimulated with 2.5 ng/mL TGF-β1, the most potent myofibroblast inducer ^[14,15]^ frequently upregulated in NSCLC at similar concentrations ^[16]^. CFs and CAFs were obtained from surgical NSCLC patients (10 LUAD, 15 LUSC). Due to their limited lifespan and availability, each assay used the patient-derived fibroblasts available at the time, resulting in partially overlapping cohorts detailed in figure legends and Table S1. To establish a mechanical benchmark of canonical pulmonary myofibroblasts as a reference for myCAFs, we examined how prolonged (≥ 3 days) TGF-β1 stimulation affects α-SMA expression/organization and cell nanomechanics in CFs. Single-cell immunofluorescence analysis based on CellMask segmentation (**Figure** 1a) showed that TGF-β1 stimulation of CFs (patient #5, where the number after # denotes the patient identifier) markedly increased α-SMA staining, largely within stress fibers (**Figure** 1b; **Figure S**2), making α-SMA^+^ cells the dominant population (henceforth referred to as myofibroblasts) (**Figure** 1b,c). TGF-β1 also shifted cells from a highly elongated, spindle-shaped morphology to a more spread-out and polygonal one (**Figure 1b; Figure S**2).

**Figure 1.**
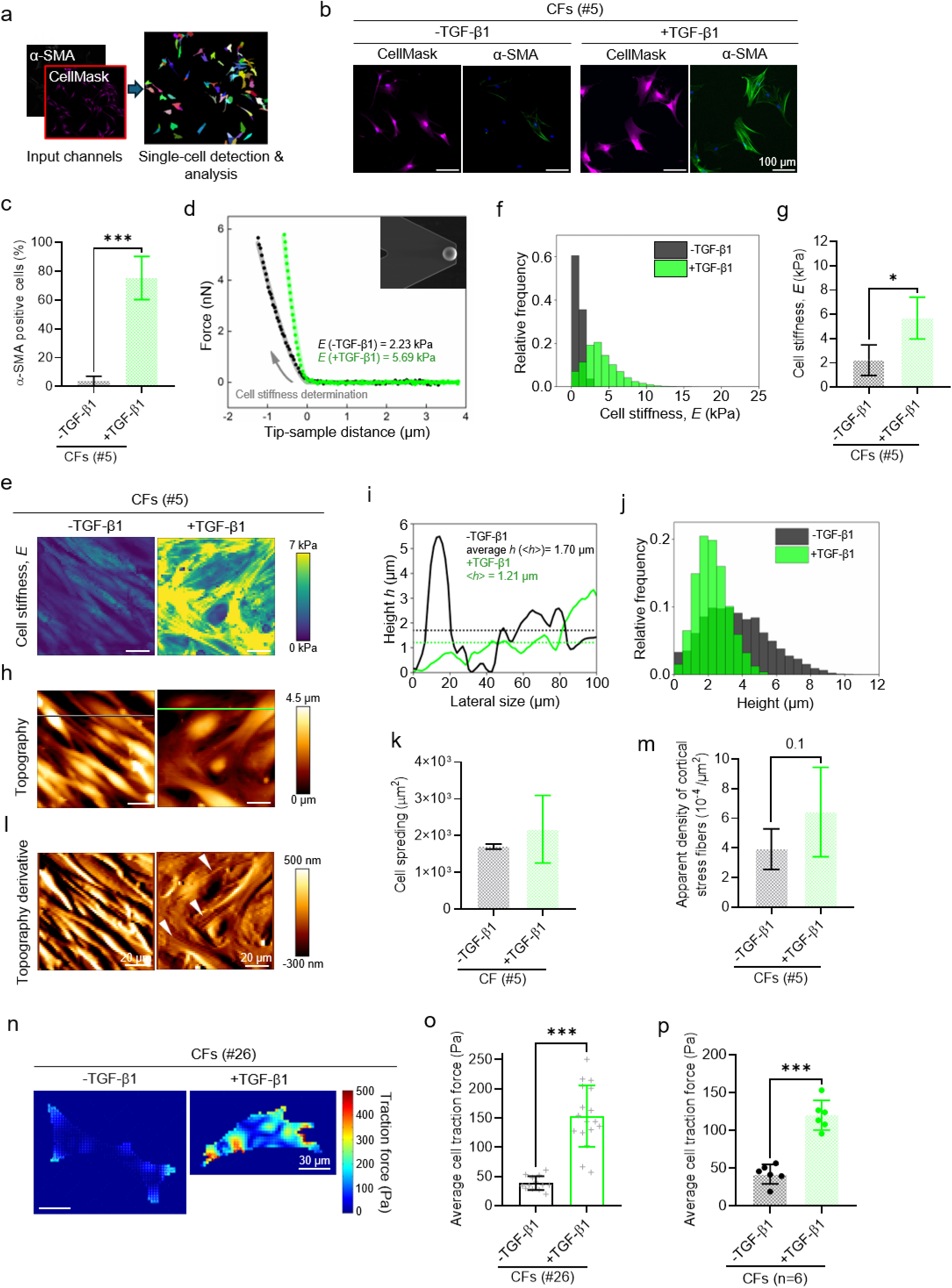
Combined AFM and TFM nanomechanical benchmarking of canonical pulmonary myofibroblast activation by TGF-β1. (a) Schematic of the CellMask-based single-cell segmentation and immunofluorescence analysis. (b) Selected regions from representative fluorescence images of CellMask and α-SMA merged with Hoechst 33342 staining of control pulmonary fibroblasts (CFs) derived from uninvolved pulmonary tissues obtained from surgical NSCLC patients (patient #5) cultured in the absence (-TGF-β1, our controls) or presence of TGF-β1 (+TGF-β1) for 5 days. Full-field images and images from additional representative regions shown in **Figure** S2. (c) Percentage (%) of α-SMA+ CFs (#5) cultured as in (b). (d) Representative AFM force-distance curves (approaching arm) acquired in a single CFs (#5) cultured in the absence or presence of TGF-β1 for 3 days at a similar height. Top-right inset shows a SEM image of a representative microspherical tip used in AFM measurements. (e) Representative maps of the Younǵs modulus (*E*) measured with AFM in CFs (#5) cultured as in (d). (f) Distributions of *E* values measured in CFs (#5) cultured as in (d) from 9 regions per sample; *E* data were filtered to correct for potential artifacts due to the underlying stiff substratum as outlined in **Figure** S3. A range of 40 to 100 fibroblasts were examined in AFM measurements per condition hereafter. (g) Average *E* values measured in CFs (#5) cultured as in (d), computed from an average number of 25932 distinct cell locations probed per sample. (h) Topography images corresponding to the stiffness maps shown in (e). (i) Height (*h*) profiles along the horizontal lines indicated in h). Dashed lines indicate the average height across each line in (h). (j) Distributions of height values from 9 regions per sample obtained in CFs (#5) cultured as in (d). (k) Singe-cell spreading assessed from CellMask segmentations in CFs (#5) cultured as in (b). (l) First derivative of the topography images shown in (h). (m) Apparent density of cortical stress fibers assessed from the first derivative of the topography images of CFs (#5) cultured as in (d), averaged from 9 images per sample. (n) Representative maps of traction forces obtained with TFM on a single CF (#26) cultured as in (b) on 6 kPa collagen-coated polyacrylamide gels. Maps from another patient shown in **Figure** S4. (o) Average single-cell traction forces of CFs (#26) cultured as in (n). Each grey symbol “+” represents an individual cell. (p) Average cell traction forces of CFs cultured as in (b) (n=6 patients, 10 cells/patient; #15, #20, #22, #26, #27, #33). Error bars represent mean ± SD of two technical replicates (g, m). Each dot corresponds to the average of a different patient (p). * *p* < 0.05; **, *p* < 0.01; ***, *p* < 0.001 with respect to control conditions (absence of exogenous TGF-β1). Statistical comparisons were done using two-tailed Student’s *t*-test.

Representative AFM force-distance curves (approach arms) acquired with a microspherical tip without (black) or with (green) TGF-β1 on CFs (#5) revealed a higher slope in the contact region (tip-sample distance < 0) with TGF-β1, indicating increased local stiffness, as confirmed by the Younǵs modulus (*E*) (**Figure** 1d). Pseudocolored stiffness (*E*) maps computed from force-distance curves (**Figure S**3) and their distributions shifted markedly towards higher stiffness with TGF-β1 (**Figure** 1e,f), with an average *E* of ∼6 kPa, three-fold higher than unstimulated controls (**Figure** 1g). The relative variability of *E* values, assessed by the coefficient of variation (SD/mean), was lower with TGF-β1 (**Figure S**3), consistent with reduced heterogeneity through a robust myofibroblast phenotype induction. AFM topography revealed a global cell height reduction with TGF-β1 (**Figure** 1h-j), together with higher cell spreading from CellMask segmentations (**Figure** 1k), in agreement with reports of TGF-β1-induced fibroblast flattening ^[35]^. The apparent density of stiff cortical stress fibers, quantified by counting long bundles along the major axis in the spatial derivative of topography maps (**Figure** 1l, white arrows), also increased with TGF-β1 with marginal significance (**Figure** 1m), consistent with their prominent mechanical role ^[26,27,32]^.

In line with AFM, traction-force maps of a single CF (#26 in **Figure** 1n; other patients in **Figure S**4) showed enhanced traction forces upon TGF-β1, particularly at the cell periphery, where focal adhesions typically develop ^[26]^. Consistently, most single-cell TFM measurements on CFs (#26) shifted toward higher traction forces with TGF-β1 (**Figure** 1o). On average, TGF-β1 raised traction forces in CFs (n=6) to 120 Pa, nearly three-fold higher than controls (**Figure** 1p; each dot represents the average for a single patient hereafter), quantitatively matching the AFM stiffness increase (**Figure** 1g). This concordance between AFM and TFM supports the robustness of our findings, indicating that TGF-β1 drives nanomechanical changes in CFs independently of inter-patient variability through coordinated increases in intracellular stiffness (AFM), extracellular force generation and transmission (TFM), and stress-fiber assembly rich in α-SMA, together with cell flattening. These nanomechanical signatures of canonical pulmonary myofibroblasts will serve as a reference for comparisons with lung myCAFs.

### 2.2. Combining AFM and TFM unveils a non-canonical hypercontractile phenotype in LUSC-myCAFs

Next, we characterized the α-SMA expression and nanomechanics of TGF-β1-stimulated lung CAFs of LUAD and LUSC origin (blue and red, respectively, hereafter). α-SMA staining showed prominent stress fibers (**Figure** 2a**; Figure S**5) and a large percentage (%) of α-SMA^+^ CAFs across histotypes (3 LUAD, 3 LUSC), moderately higher in LUAD (**Figure** 2b). On average, the % of α-SMA^+^ CAFs was comparable to that of CFs with TGF-β1 (**Figure** 1c), confirming a dominant myCAF population. Accordingly, TGF-β1-treated CAFs were termed myCAFs. As in CFs, TGF-β1 elicited a spread-out polygonal shape in myCAFs, yet the surface area was significantly lower in LUSC (**Figure** 2a,c; **Figure** S5). AFM measurements on myCAFs (4 LUAD, 4 LUSC) were performed in LUAD-LUSC histotype pairs (**Figure** S6) at high tumor-like cell densities, as in CFs (**Figure S**1). Force-distance curves from a pairwise comparison revealed a steeper slope in LUSC-myCAFs (#16) than LUAD-myCAFs (#7), indicating higher local stiffness in LUSC (**Figure** 2d). Consistently, *E* maps (**Figure** 2e,f, left panels) and their distributions (**Figure** 2g; **Figure S**6) shifted towards higher values in LUSC-myCAFs in all pairwise comparisons, with an average *E* 70% higher than LUAD-myCAFs (**Figure** 2h). This finding was unexpected, considering the prominent α-SMA expression in both histotypes and the modestly lower % of α-SMA^+^ cells in LUSC (**Figure** 2b). Conversely, the coefficient of variation of *E* was consistently lower in LUSC-myCAFs (Figure S7), revealing a more uniform nanomechanical phenotype. As in the coordinated stiffening and flattening of canonical pulmonary myofibroblasts (**Figure** 1g-j), height distributions from topography images (**Figure** 2e,f, middle panels) shifted toward lower values in LUSC-myCAFs (**Figure** 2i,j); together with reduced spreading (**Figure** 2c), this indicates a smaller cell size in LUSC-myCAFs. Moreover, the apparent density of stiff stress fibers assessed from the first derivative of topography images (**Figure** 2e,f, right panels; **Figure** S6) was significantly higher in LUSC-myCAFs (**Figure** 2k), consistent with their relevant mechanical role.

**Figure 2.**
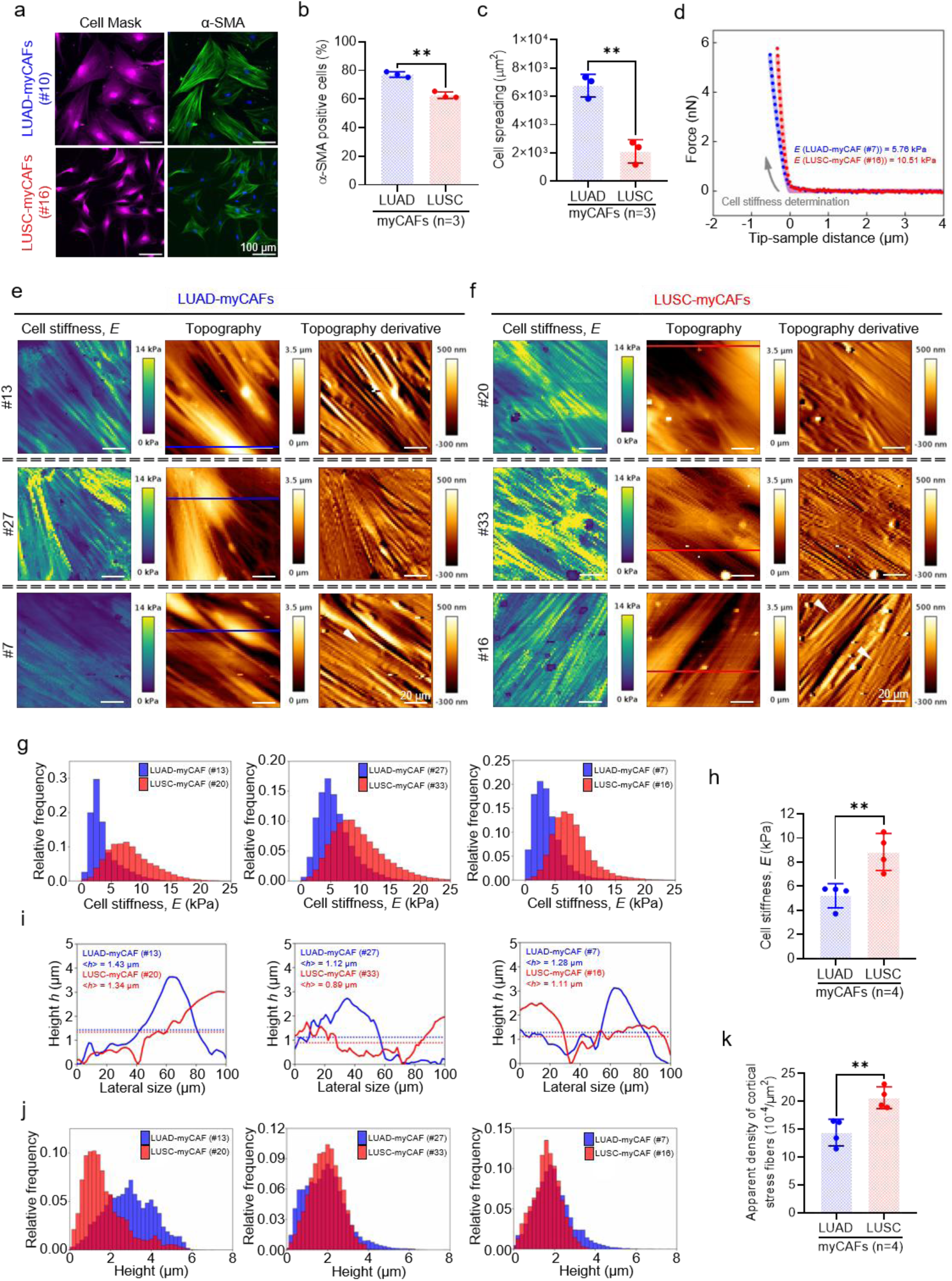
AFM reveals histotype-dependent nanomechanical phenotypes in lung myCAFs. (a) Selected regions from representative images of CellMask and α-SMA merged with Hoechst 33342 nuclear staining of cancer-associated fibroblasts (CAFs) obtained from surgical NSCLC patients (LUAD and LUSC) and stimulated with TGF-β1 for 5 days to induce a robust myofibroblast-like phenotype (henceforth referred to as myCAFs). Full-field images and images from additional fields and patients shown in **Figure** S5. (b) Percentage (%) of α-SMA+ myCAFs (3 LUAD, 3 LUSC; #4, #6, #7, #10, #12, #16) cultured as in (a). (c) Singe-cell spreading area of the myCAFs in (b), assessed from CellMask segmentations (average cell number = 847 for LUAD, 3655 for LUSC). (d) Representative AFM force-distance curves (approaching arm) acquired in single myCAFs from LUAD and LUSC patients stimulated with TGF-β1 for 3 days at a similar height. (e-f) Representative AFM maps of cell stiffness (left), topography (middle), and topography first derivative (right) for LUAD-myCAFs (e) and LUSC-myCAFs (f) cultured as in (d). Each row shows measurements acquired from a LUAD-LUSC histotype pair within the same experiment (all pairs listed in **Figure** S6). (g) Distributions of *E* values of myCAFs cultured as in (d) from each histotype pairwise measurement shown in (e-f) (9 regions per sample). *E* data were processed as in **Figure** 1f. (h) Average *E* values measured in myCAFs (4 LUAD, 4 LUSC) cultured as in (d), computed from an average number of 48128 distinct cell locations per patient sample. (i) Height (*h*) profiles along the horizontal lines indicated in the topography images in e-f). Dashed lines indicate the corresponding average height (*h*) across the profile. (j) Distributions of height values of myCAFs cultured as in (d) (9 images/sample) from the histotype pairs shown in (e-f). (k) Apparent density of cortical stress fibers assessed from the first derivative of the topography images of myCAFs (4 LUAD, 4 LUSC; 9 images/sample) cultured as in (d). Error bars represent mean ± SD of two technical replicates (h, k). Each dot corresponds to the average of a different patient (b, c, h, k). **, *p* < 0.01 comparing LUAD-myCAFs and LUSC-myCAFs. Statistical comparisons were done using two-tailed Student’s *t*-test.

As in canonical pulmonary myofibroblasts (**Figure** 1n), high traction forces in single LUAD- and LUSC-myCAFs localized largely at the cell periphery (**Figure** 3a). In line with AFM results, LUSC-myCAFs exhibited higher single-cell average traction forces than LUAD-myCAFs (3 LUAD, 3 LUSC) (**Figure** 3b), which were collectively 40% larger (**Figure** 3c), and a lower coefficient of variation of the average traction forces per patient (**Figure S**7), confirming a more homogeneous nanomechanical phenotype. The hypercontractile phenotype of LUSC-myCAFs, independently observed by AFM and TFM in partially overlapping patient cohorts, supports the robustness of our approach, and confirms that histotype-dependent differences exceed inter-patient variability. Unstimulated LUSC-CAFs also showed significantly higher traction forces than LUAD-CAFs despite similar α-SMA staining (**Figure S**8), indicating that the histotype nanomechanical divergence is intrinsic rather than dependent on exogenous TGF-β1. Notably, stiffness (**Figure** 2h), traction forces (**Figure** 3c) and stress fiber density (**Figure** 2k) in LUAD-myCAFs closely matched those of TGF-β1-stimulated CFs (**Figure** 1g,p,m, respectively), indicating that LUAD-myCAFs retain the phenotype of canonical pulmonary myofibroblasts. In contrast, LUSC-myCAFs were even more contractile through a coordinated amplification of stiffness, traction-force generation and transmission, and stress-fiber density, concomitantly with enhanced cell size reduction, unveiling a previously unrecognized non-canonical hypercontractile phenotype.

**Figure 3.**
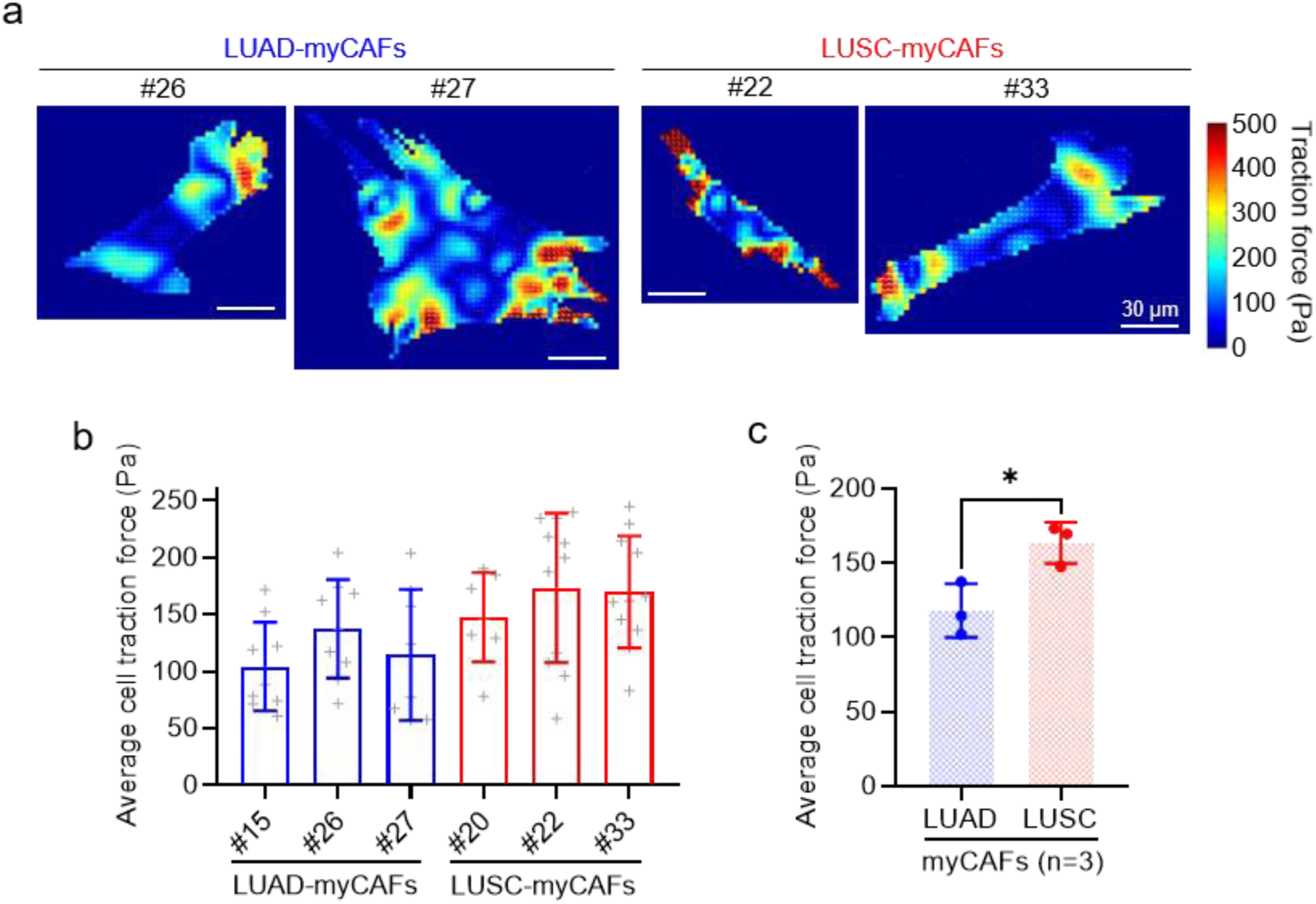
AFM confirms a non-canonical hypercontractile phenotype in LUSC-myCAFs. (a) Representative maps of traction forces obtained with TFM on single LUAD-myCAFs (left panels) and LUSC-myCAFs (right panels) stimulated with TGF-β1 for 5 days. (b) Average single-cell traction forces of myCAFs from individual patients (3 LUAD, 3 LUSC) cultured as in a). Each grey symbol “+” represents an individual cell (9 cells/patient in average). (c) Average cell traction forces of myCAFs shown in (b). Each dot represents a single patient (3 LUAD, 3 LUSC). Error bars represent mean ± SD. * *p* < 0.05; **, *p* < 0.01; ***, *p* < 0.001 comparing LUAD-myCAFs and LUSC-myCAFs. Statistical comparisons were done using two-tailed Student’s *t*-test.

### 2.3. LUSC-myCAFs exhibit enhanced smooth muscle myosin heavy chain (*MYH11*) expression and colocalization with the α-SMA cytoskeleton, consistent with the acquisition of a smooth muscle cell (SMC)-like contractile program

Prompted by the histotype-dependent nanomechanics of lung myCAFs, we investigated their molecular basis. Because α-SMA is widely regarded as the main driver of canonical myofibroblast contractility through its interaction with NMMII motors ^[15,22]^, we first systematically investigated whether differences in α-SMA expression could account for the hypercontractility of LUSC-myCAFs, using different techniques and multiple cohorts. All fibroblasts were stimulated with TGF-β1 as in nanomechanical measurements. *ACTA2* (α-SMA gene) RNA levels were significantly higher in LUAD-myCAFs than LUSC-myCAFs (8 LUAD, 5 LUSC) (**Figure** 4a), whereas α-SMA protein levels were similar in both histotypes by Western blotting (WB) (4 LUAD, 4 LUSC) (**Figure** 4b) and immunofluorescence (3 LUAD, 3 LUSC) (**Figure** 4c). Likewise, *ACTA2* levels in bulk tumor transcriptomes from the Tissue Cancer Genome Atlas (TCGA) (706 LUAD, 626 LUSC) were significantly higher in LUAD (**Figure** 4d), but this difference did not hold at the tissue level, as the % of α-SMA^+^ stromal area was comparable across histotypes in tissue microarrays (TMAs) from the CIBERES multicenter Spanish cohort (105 LUAD, 92 LUSC) ^[8]^ (**Figure** 4e,f) and the Human Protein Atlas (HPA) (**Figure** 4g,h). Thus, α-SMA (*ACTA2*) expression alone is insufficient to explain the hypercontractile phenotype of LUSC-myCAFs, suggesting alternative contractility drivers.

**Figure 4.**
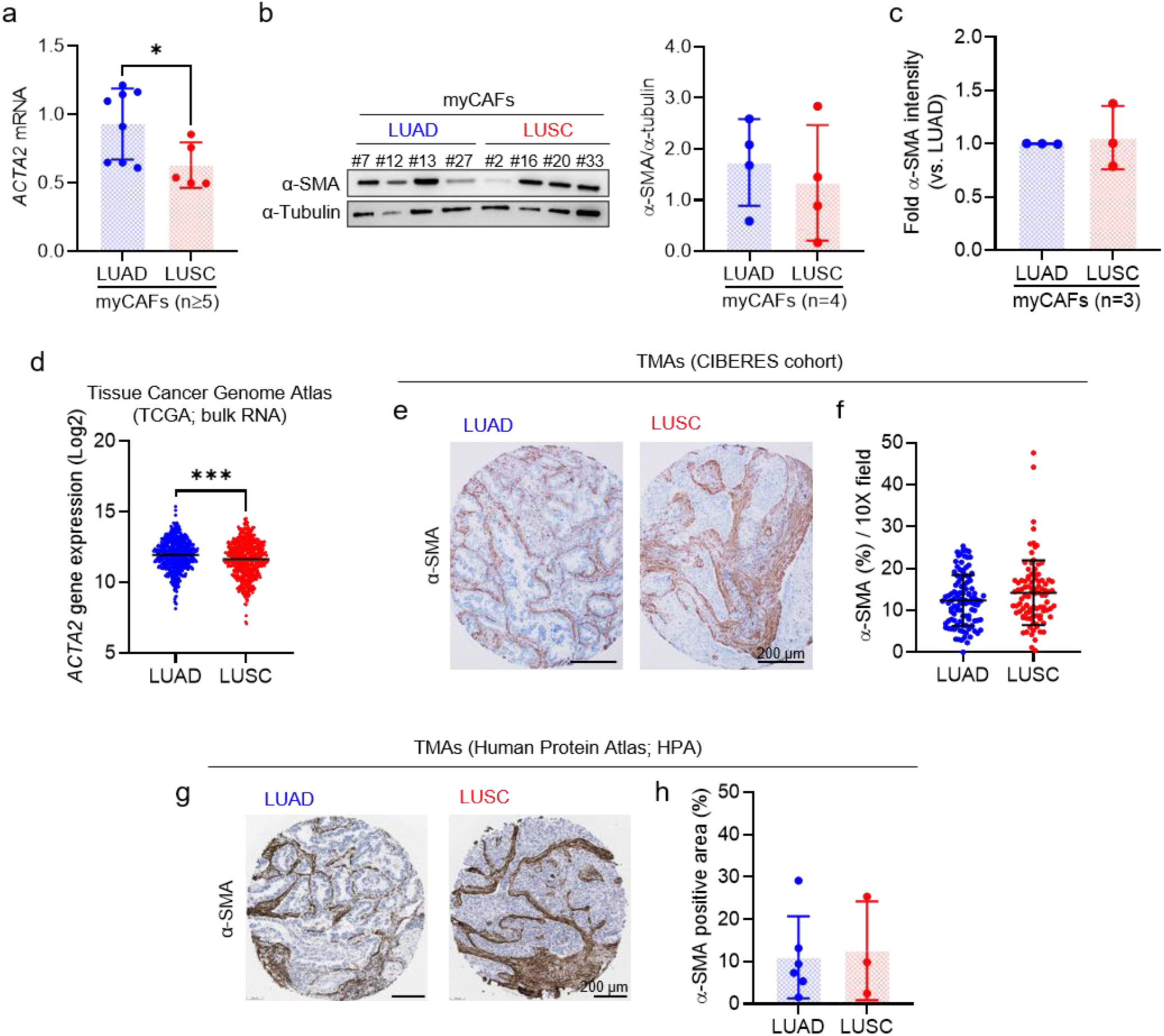
Histotype-Specific myCAF nanomechanics are not explained by α-SMA (*ACTA2*) expression. (a) mRNA levels of *ACTA2* (α-SMA) assessed by qRT-PCR (three technical replicates) on myCAFs stimulated with TGF-β1 for 3 days (8 LUAD, 5 LUSC). (b) (left) Western blot of α-SMA and α-Tubulin in myCAFs cultured as in (a); (right) corresponding densitometry analysis of α-SMA/α-Tubulin. (c) Single-cell α-SMA immunofluorescence intensity in myCAFs cultured as in **Figure** 2a (3 LUAD, 3 LUSC), normalized to the mean value of LUAD-myCAFs. (d) Tumor bulk *ACTA2* (α-SMA) mRNA levels in TCGA NSCLC cohorts (706 LUAD, 626 LUSC). (e) Representative images of histologic staining of α-SMA in TMAs from LUAD and LUSC patients within the CIBERES cohort. (f) Average quantification of the percentage (%) of positively stained α-SMA area per patient in TMAs from the CIBERES cohort (105 LUAD, 92 LUSC). (g) Representative images of α-SMA histological staining in TMAs from LUAD and LUSC tumors obtained from the Human Protein Atlas (HPA) database (antibody CAB000002). (h) Average quantification of the percentage (%) of positively stained α-SMA area per patient in TMAs from the HPA cohort (6 LUAD, 3 LUSC). Error bars represent mean ± SD. Each dot corresponds to the average of a different patient (a-h). * *p* < 0.05; **, *p* < 0.01; ***, *p* < 0.001 comparing LUAD and LUSC. Statistical comparisons were done using two-tailed Student’s *t*-test.

A previous spatial transcriptomic and proteomic study identified an abundant NSCLC myCAF population co-expressing α-SMA and smooth muscle myosin heavy chain (SM-MHC, encoded by *MYH11*), referred to as α-SMA^+^/SM-MHC^+^, together with overexpression of contractility-related genes ^[11]^. SM-MHC is the core component of smooth muscle myosin II (SMMII) motors, which drive bulk force generation in smooth muscle cells (SMCs), largely through the α-SMA cytoskeleton ^[38]^, and SMMII has a greater force-generating capacity than NMMII ^[39]^. Although SM-MHC is traditionally considered low or absent in canonical myofibroblasts ^[15]^, its contractile role and previous observations in NSCLC motivated us to examine the histotype-specific co-expression of α-SMA and SM-MHC in myCAFs by dual immunofluorescence (3 LUAD, 3 LUSC). SM-MHC displayed diffuse cytoplasmic distribution (**Figure** 5a, first column; **Figure** S9), as previously reported ^[40]^, unlike the stress-fiber organization of α-SMA (**Figure** 5a, second column; **Figure** S9).

**Figure 5.**
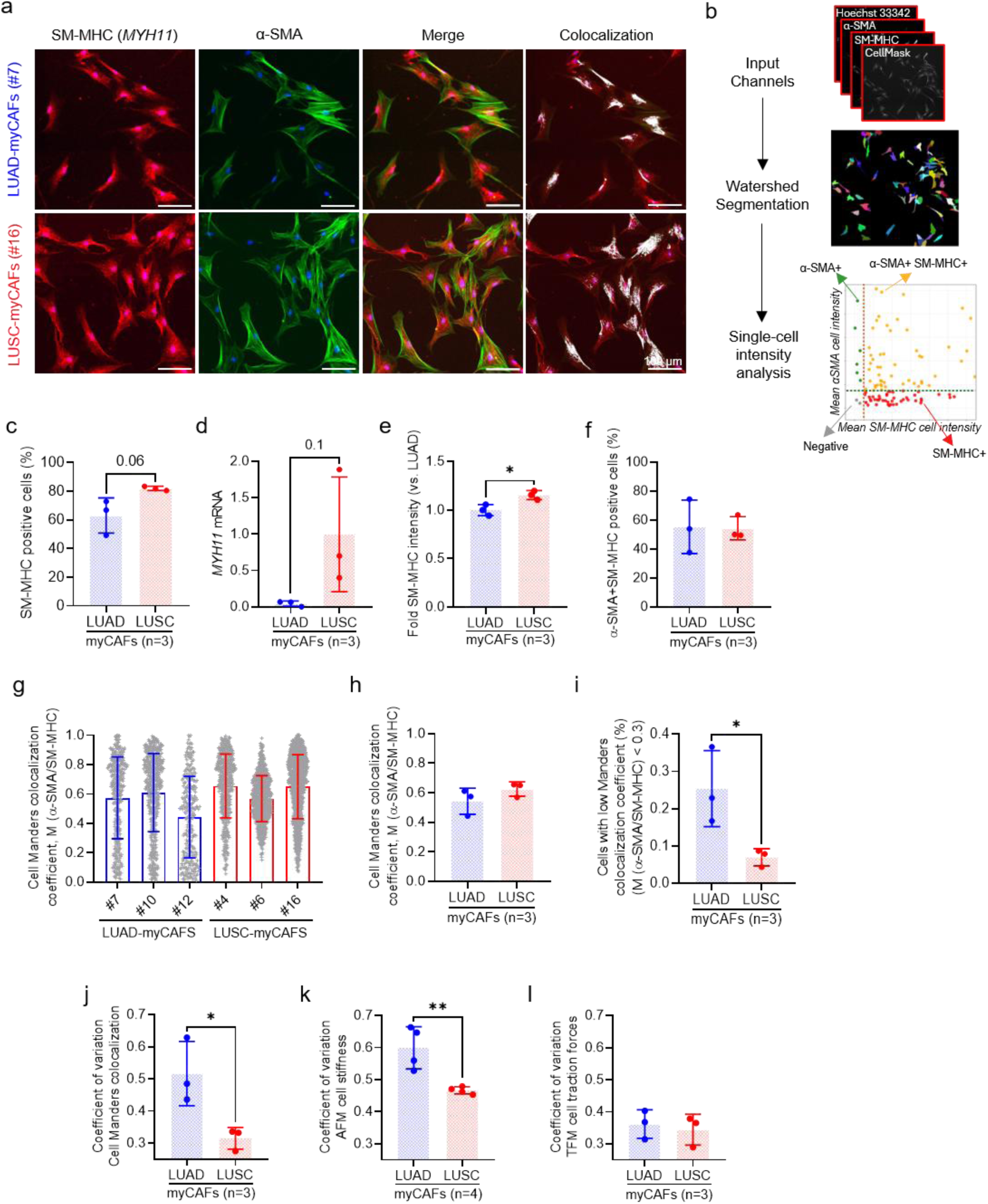
Enhanced smooth muscle myosin heavy chain (SM-MHC (*MYH11*)) expression and incorporation into the α-SMA cytoskeleton support an smooth muscle cell (SMC)-like phenotype in LUSC-myCAFs. (a) Selected regions from representative images of SM-MHC (*MYH11*) (first column) and α-SMA (second column) immunofluorescence co-staining in LUAD-myCAFs (#7) and LUSC-myCAFs (#16) cultured as in **Figure** 2a. Images of the merged channels and their colocalization (labelled in white) are shown in the third and fourth columns, respectively. Nuclei were counterstained with Hoechst 33342. Full-field images and images from additional fields and patients shown in **Figure** S9. (b) Schematic of the CellMask-based single-cell analysis of the co-immunofluorescence staining for SM-MHC and α-SMA. Additional details in **Figure** S10. (c) Percentage (%) of SM-MHC+ myCAFs (3 LUAD, 3 LUSC; #4, #6, #7, #10, #12, #16) cultured as in (a). (d) mRNA levels of *MYH11* assessed by qRT-PCR (three technical replicates) on myCAFs (3 LUAD, 3 LUSC; #4, #6, #7, #12, #13, #20) cultured as in (a). (e) Single-cell SM-MHC (*MYH11*) immunofluorescence intensity in myCAFs cultured as in **Figure** 2a, normalized to the mean value of LUAD-myCAFs (3 LUAD, 3LUSC). (f) Percentage (%) of double positive (α-SMA+ SM-MHC+) myCAFs (3 LUAD, 3LUSC) cultured as in (a). (g) Single-cell Manders colocalization coefficient M(α-SMA/SM-MHC) quantifying the fraction of α-SMA co-occurring with SM-MHC normalized by total α-SMA in myCAFs from individual patients (3 LUAD, 3 LUSC) cultured as in (a). Each grey symbol “+” represents an individual cell (average cell number analyzed/patient: 617). (h) Average cell Manders colocalization coefficient M(α-SMA/SM-MHC) corresponding to data shown in (g). (i) Average percentage (%) of cells in (g) with low Manders colocalization coefficient (M(α-SMA/SM-MHC) < 0.3). Similar analysis considering M(α-SMA/SM-MHC) < 0.2 shown in **Figure** S11. (j) Average coefficient of variation (SD/mean) of cell Manders colocalization data shown in (g). (k) Average coefficient of variation of AFM cell stiffness data shown in **Figure** 2j and **Figure** S6. (l) Average coefficient of variation of TFM data shown in **Figure** 3b. Each dot corresponds to the average of a different patient (c-f, h-l). Error bars represent mean ± SD. * *p* < 0.05; **, *p* < 0.01; ***, *p* < 0.001 comparing LUAD and LUSC. Statistical comparisons were done using two-tailed Student’s *t*-test.

CellMask based single-cell analyses (**Figure** 5b; **Figure** S10) showed that SM-MHC^+^ myCAFs were abundant in both histotypes, yet more frequent in LUSC (∼80%) than LUAD (∼60%) with marginal significance (3 LUAD, 3 LUSC) (**Figure** 5c). Consistently, *MYH11* RNA (**Figure** 5d) and SM-MHC fluorescence intensity (**Figure** 5e) were significantly higher in LUSC-myCAFs, even in basal conditions (**Figure** S8), supporting an intrinsic priming towards an SMC-like contractile phenotype. In contrast, the % of α-SMA^+^/SM-MHC^+^ cells was similar between histotypes (Figure 5A, third column; **Figure** 5f), indicating that differences in contractility could not be explained by selective enrichment of double-positive cells. We therefore assessed the integration of SM-MHC within the α-SMA cytoskeleton (**Figure** 5a fourth column) using the Manders colocalization coefficient M(α-SMA/SM-MHC), which quantifies the fraction of α-SMA co-occurring with SM-MHC over total α-SMA within individual cells, and ranges from 0 (absence) to 1 (complete colocalization) (**Figure** 5g). The mean colocalization coefficient was only modestly higher in LUSC-myCAFs (**Figure** 5h), but the distributions differed markedly, with substantially greater variability in LUAD-myCAFs spanning the low colocalization range (**Figure** 5g). Accordingly, a significantly larger fraction of cells with very low colocalization (M < 0.3) was found in LUAD-myCAFs (**Figure** 5i); consistent results were obtained for M < 0.2 (**Figure** S11). The more compact LUSC distributions yielded a lower coefficient of variation (**Figure** 5j), as observed by AFM and TFM (**Figure** 5k,l; **Figure** S7), indicating that LUSC-myCAFs adopt a more uniform contractile phenotype. The closer resemblance of LUSC-myCAFs to SMCs in terms of enhanced SM-MHC (*MYH11*) expression and its incorporation into the α-SMA cytoskeleton supports the involvement of SMMII in their hypercontractility.

### 2.4. Contractility-regulatory signaling, DNA methylation reprogramming, and stromal enrichment of SMC markers reveal a strong activation of an SMC-like contractile program in LUSC-myCAFs

To further assess the mechanical convergence between LUSC-myCAFs and SMCs, we analyzed major upstream regulators of NMMII and SMMII, the key contractility effectors in canonical myofibroblasts and SMCs, respectively ^[15,41]^. Although both NMMII and SMMII are activated through regulatory light chain phosphorylation (RLC^pS19/S20^) ^[42]^, the contributions of their two major upstream kinases, ROCK and myosin light-chain kinase (MLCK), differ. Thus, canonical myofibroblasts rely substantially on ROCK-mediated inhibition of myosin light-chain phosphatase (MLCP) via phosphorylation of its regulatory subunit MYPT1 (MYPT1^pT853^) ^[14,28,43]^, whereas SMC contractility depends more heavily on MLCK, which directly activates SMMII ^[41,44–46]^, and can be further enhanced by ERK1/2 phosphorylation (ERK1/2^pT202/Y204^) ^[47,48]^. WB of TGF-β1-stimulated CAFs showed significantly lower MYPT1^pT853^ (**Figure** 6a,b) and higher ERK1/2^pT202/Y204^ (**Figure** 6c,d) in LUSC-myCAFs than LUAD-myCAFs (≥3 LUAD, 4 LUSC), revealing a shift from canonical ROCK-MYPT1 regulation toward SMC-like signaling in LUSC-myCAFs. RLC^pS19/S20^ was also higher in LUSC-myCAFs (**Figure** 6a, e), although the difference was smaller than that observed for MYPT1^pT853^ and ERK1/2^pT202/Y204^, and was not significant. However, because SMMII generates higher forces than NMMII ^[39,49]^, a moderate increase in RLC phosphorylation could translate into substantially greater contractility if part of the phosphorylated RLC is associated with SMMII, further supporting SMMII involvement in LUSC-myCAF nanomechanics.

**Figure 6.**
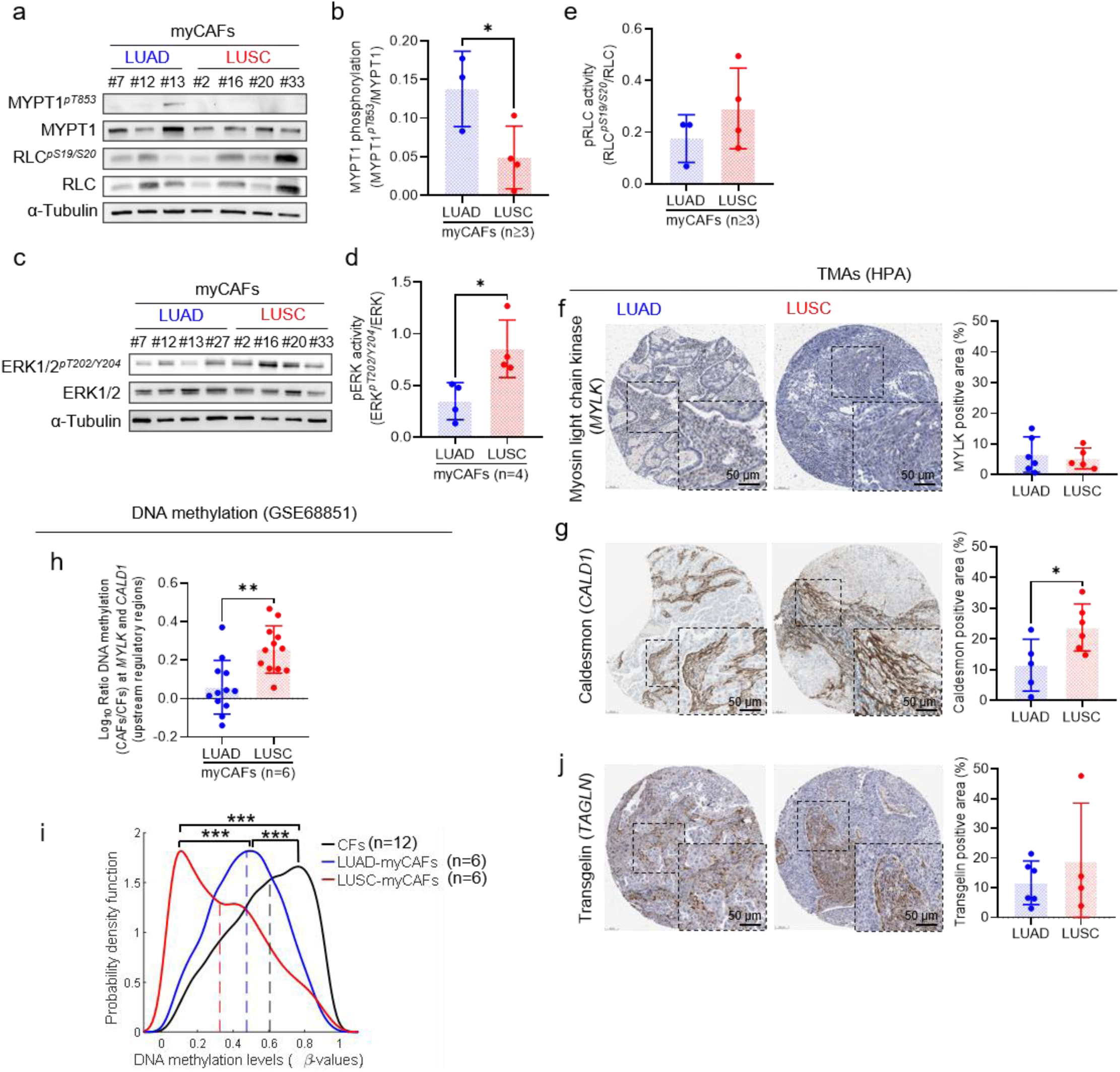
Signaling and epigenetic evidence for an SMC-Like contractile program in LUSC-myCAFs. (a) Western blot of MYPT1^pT853^, total MYPT1, RLC^pS19/S20^, total RLC, and α-Tubulin of myCAFs (3 LUAD, 4 LUSC) stimulated with TGF-β1 for 20 min. (b) Densitometry analysis of MYPT1^pT853^/total MYPT1 from bands shown in (a). (c) Western blot of ERK1/2^pT202/Y204^, total ERK1/2, and α-Tubulin of myCAFs (3 LUAD, 4 LUSC) cultured as in (a). (d) Densitometry analysis of ERK1/2^pT202/Y204^/total ERK from bands shown in (c). (e) Densitometry analysis of RLC^pS19/S20^/total RLC from bands shown in (a). (f) (left) Representative images of myosin light-chain kinase (encoded by *MYLK*) histological staining in TMAs from LUAD and LUSC tumors obtained from the HPA (antibody CAB020789). Inset shows a magnified view of the indicated region; (right) corresponding average quantification of the percentage (%) of positively stained area per patient (6 LUAD, 5 LUSC). (g) (left) Representative images of caldesmon (encoded by *CALD1*) histological staining in TMAs from LUAD and LUSC tumors obtained from the HPA (antibody HPA008066). Inset shows a magnified view of the indicated region; (right) corresponding average quantification of the percentage (%) of positively stained area per patient (5 LUAD, 6 LUSC). (h) Log10-transformed ratio (CAFs relative to paired CFs) of the average DNA methylation level (β-value, from GSE68851 dataset) for selected CpG sites located in the proximal (*MYLK*) or distal (*CALD1*) regulatory regions (CpG sites detailed in the Experimental Section). Values below 0 indicate lower methylation in CAFs. Each dot represents the score for one gene in an individual patient (6 LUAD, 6 LUSC: #5, #6, #9, #12, #13, #14, #16, #18, #25, #26, #27, #28). Cells were cultured in basal conditions (without exogenous TGF-β1). (i) Distribution of DNA methylation levels (β-values, 0–1) of the 1452 CpG sites with the largest differential methylation between CAFs and CFs (GSE68851 dataset; 12 CFs, 6 LUAD-CAFs, 6 LUSC-CAFs). Dashed vertical lines indicate median β-values of each group. (j) (left) Representative images of transgelin (encoded by *TAGLN*) histological staining in TMAs from LUAD and LUSC tumors obtained from the HPA (antibody CAB001447). Inset shows a magnified view of the indicated region; (right) corresponding average quantification of the percentage (%) of positively stained area per patient (6 LUAD, 4 LUSC). Each dot corresponds to the average of a different patient (b, d-h, j). Error bars represent mean ± SD. * *p* < 0.05; **, *p* < 0.01; ***, *p* < 0.001 comparing LUAD and LUSC. Statistical comparisons were done using two-tailed Student’s *t*-test. Wilcoxon rank sum test was used in (i).

To independently support the selective enrichment of an SMC-like contractile program in LUSC-myCAFs, we benefited from our previous DNA methylation profiling of lung CAFs and paired CFs, as it reported a global enrichment of the KEGG vascular smooth muscle contraction pathway (hsa04270) in CAFs ^[36]^, although histotype-dependent patterns were not considered. Reanalysis of this dataset (GSE68851; 6 LUAD and 6 LUSC) revealed that, among the four direct actomyosin regulators in this pathway with stronger epigenetic remodeling in CAFs than CFs (*MYLK*, *CALD1*, *GNA12* and *PRKCE*), only the expression of *MYLK* and *CALD1* (encoding for MLCK and caldesmon, respectively) was largely restricted to the stroma in HPA TMAs (**Figure** 6f,g; **Figure** S12). Systematic analysis of differential DNA methylation levels between CAFs and CFs within main promoter regions and upstream regulatory sites in *MYLK* and *CALD1* showed significantly larger alterations in LUSC-CAFs than LUAD-CAFs in CpG sites within upstream regulatory regions (proximal for *MYLK*, distal for *CALD1*) (**Figure** 6h), indicating larger epigenetic modifications within alternative transcription start sites. Likewise, the global DNA hypomethylation of CAFs relative to CFs that we previously reported ^[36]^ was more pronounced in LUSC, with lower median β-values than LUAD-CAFs and CFs (horizontal dashed lines, **Figure** 6i). Collectively, these results indicate broader epigenetic remodeling in LUSC-CAFs, including selective modifications in major SMC contractility regulatory genes (*MYLK* and *CALD1*). To validate our observations at the tissue level, we examined additional canonical SMC markers, SM-MHC (*MYH11*), transgelin/SM22α (*TAGLN*) and smoothelin (*SMTN*) ^[15,50,51]^, in HPA TMAs. Only transgelin could be reliably assessed, showing a trend toward higher expression in LUSC that did not reach statistical significance, possibly due to sample size limitations (**Figure** 6j) (6 LUAD, 4 LUSC); the other markers showed low staining quality (**Figure** S12). Altogether, these results point to the prominent activation of an SMC-like contractile program in LUSC-myCAFs concomitantly with larger epigenetic reprogramming.

### 2.5. SMC-like myCAF features are associated with poor prognosis and distinctive stromal remodeling patterns selectively in LUSC

To assess the broader relevance of the SMC-like contractile program of LUSC-myCAFs, we first examined the prognostic value of canonical SMC markers (*MYH11*, *TAGLN*, *SMTN*, *CALD1*) within the TCGA (498 LUAD, 486 LUSC).

Notably, high expression of each marker was significantly associated with poorer survival in LUSC, whereas in LUAD, this prognostic effect was limited to *CALD1* and was inconsistent for *MYH11* (**Figure** 7a-d). Likewise, adverse outcomes correlated significantly with high *ACTA2* alone or combined with *MYH11* in LUSC but not LUAD (**Figure** S13), collectively supporting a LUSC-specific link between SMC-like features and poor outcome. Next, we explored whether the distinct contractile phenotypes of myCAFs could contribute to two well-established histotype-dependent stromal features of NSCLC: CAF density and tumor hypoxia ^[4,10,17]^. We previously reported significantly lower CAF density in LUSC than LUAD ^[10,17]^, but the mechanism remains unclear. Because cell cycle progression requires coordinated cell growth and volume expansion ^[52]^, and contractility has been reported to reduce cell volume in mesenchymal cells ^[53]^, we hypothesized that the coordinated hypercontractility and cell size reduction of LUSC-myCAFs could impair their cell cycle progression. To test this possibility, we stimulated CAFs with 10% FBS and examined the % of growth-arrested (G0/G1) cells relative to serum-deprived conditions (0% FBS) by flow cytometry. TGF-β1 was excluded, owing to its known cytostatic effects and because greater traction forces in LUSC-CAFs relative to LUAD-CAFs were also observed without TGF-β1 (**Figure S**8). In agreement with our hypothesis, LUAD-CAFs showed a reduction in growth-arrested cells upon serum stimulation (−12.45% on average, marginally significant), whereas LUSC-CAFs remained largely unresponsive (−0.15%, n.s.) (6 LUAD, 5 LUSC) (**Figure** 7e,f), consistent with intrinsic resistance to mitogenic stimulation. Next, we asked whether the greater hypoxia and necrosis reported in LUSC ^[4]^ were related to vascular compression, a recognized consequence of elevated tumor mechanical stresses such as increased stiffness ^[19]^. We assessed the blood vessel lumen size, a common proxy of tissue mechanical stress ^[19]^, on HPA histologic images of endothelial cell markers (CD31, CD34) ^[4]^ using automated morphometric analysis. Both markers consistently showed smaller vessel lumens in LUSC than LUAD with marginal significance (≥ 5 LUAD, ≥ 5 LUSC) (**Figure** 7g-j). Although these findings do not establish causality, they link the hypercontractile phenotype of LUSC-myCAFs with CAF density and hypoxia patterns observed in LUSC.

**Figure 7.**
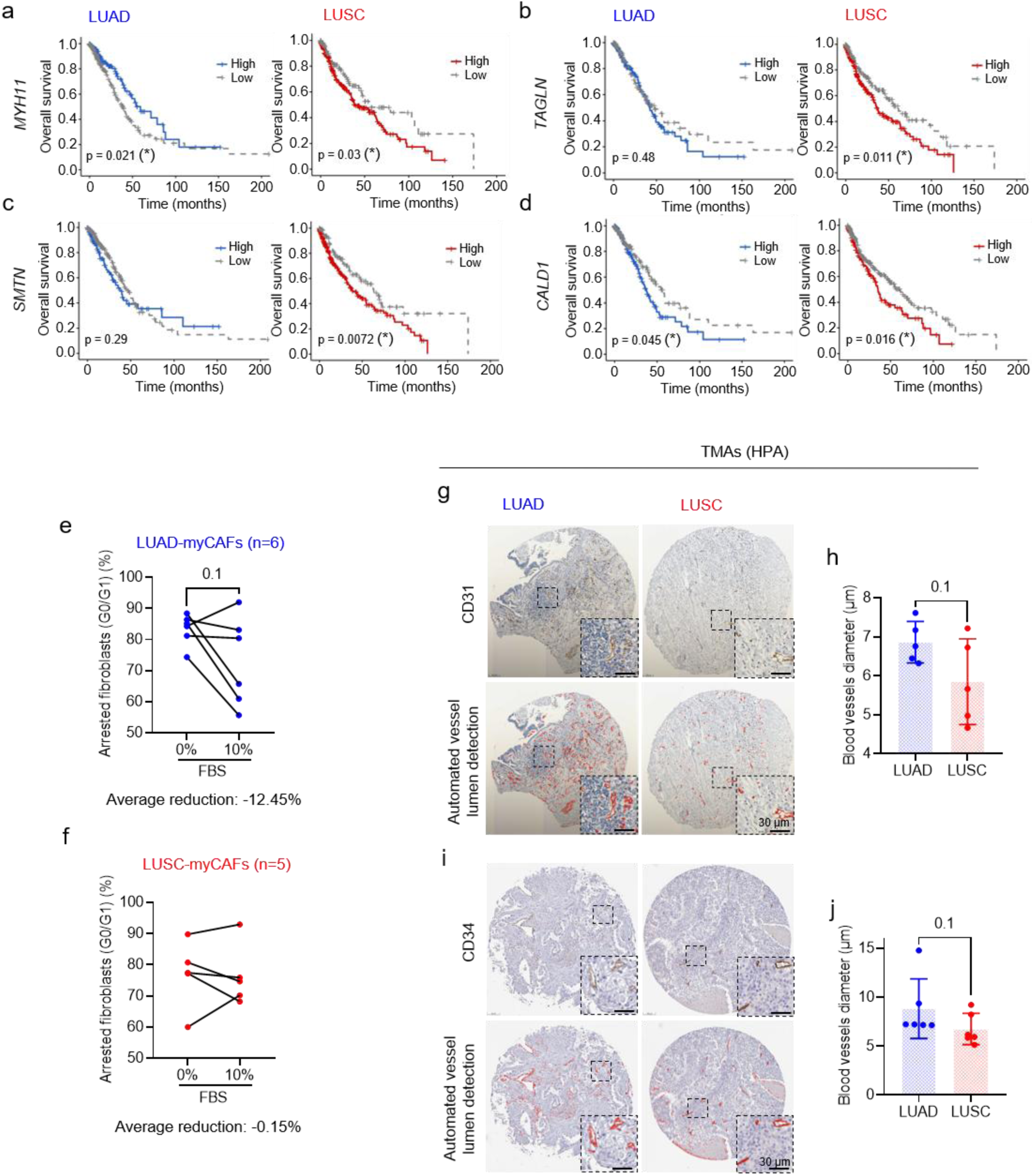
The SMC-like contractile program of LUSC-myCAFs is associated with poor prognosis and LUSC-specific patterns of stromal remodeling. (a-d) Kaplan-Meier survival curves stratifying patients according to standard SMC markers, including *MYH11* (a), *TAGLN* (b), *SMTN* (c) and *CALD1* (d). Survival curves from *ACTA2* alone or coexpressed with *MYH11* in **Figure** S13. Survival data obtained from TCGA dataset (498 LUAD, 486 LUSC). (*) indicates significant p-value. (e-f) Percentage (%) of growth arrested (G0/G1-phase) LUAD-CAFs (e) and LUSC-CAFs (f) cultured for 5 days in 0% or 10% FBS (6 LUAD, 5 LUSC; #5, #6, #9, #12, #13, #14, #15, #17, #23, #26, #28). Growth-arrested cells were determined by propidium iodide staining and flow cytometry analysis of their DNA content. The average reduction in the percentage of G0/G1 cells following serum stimulation relative to controls is shown at the bottom of each panel. (g) Representative images of CD31 histological staining (top) and corresponding automated vessel lumen detections (labelled in red, bottom panels) in TMAs from LUAD and LUSC tumors obtained from the HPA (antibody HPA004690). Insets show a magnified view of the indicated regions. (h) Average quantification of the percentage (%) of CD31 positively stained area per patient (5 LUAD, 5 LUSC). (i) (left) Representative images of CD34 histological staining (top) and corresponding automated vessel lumen detections (labelled in red, bottom panels) in TMAs from LUAD and LUSC tumors obtained from the HPA (antibody HPA036722). Insets show a magnified view of the indicated regions. (j) Average quantification of the percentage (%) of CD34 positively stained area per patient (6 LUAD, 6 LUSC). Each dot corresponds to the average of a different patient (e,f,h,j). Error bars represent mean ± SD. * *p* < 0.05; **, *p* < 0.01; ***, *p* < 0.001 comparing LUAD and LUSC. Survival curves were compared by logrank test (a-d). Statistical comparisons were done using two-tailed Student’s *t*-test or Mann-Whitney test (j).

## 3. Discussion

There is growing interest in myCAF mechanics, owing to their central role in driving the chronic tumor stiffening, a hallmark of solid tumors increasingly implicated in tumor progression and therapy resistance ^[18,19]^. Yet, studies on CAF mechanics are scarce, lack myCAF focus, and have yielded conflicting results relative to normal (control) fibroblasts, with CAFs reported as softer in pancreatic cancer ^[54]^, similarly stiff in colorectal cancer ^[34]^, or stiffer in breast and prostate cancers ^[35,55,56]^. These discrepancies likely reflect unaccounted-for fibroblast heterogeneity, potentially amplified by the common absence of exogenous TGF-β1, as CAFs frequently overexpress α-SMA relative to CFs ^[10,34,55,56]^; given that α-SMA is a TGF-β1-induced driver of myofibroblast contractility, comparisons with unstimulated CFs may confound cancer-associated differences with basal fibroblast activation levels. By combining AFM and TFM for the first time to study myCAFs, we robustly identified myCAF nanomechanics as a novel distinguishing feature between LUAD and LUSC.

While LUAD-myCAFs displayed the nanomechanical hallmarks of canonical pulmonary myofibroblasts, LUSC-myCAFs exhibited a coordinated increase in intracellular stiffness, traction-forces, and stress-fiber density beyond LUAD-myCAFs, concomitantly with a signaling shift toward contractile SMCs ^[50,51]^. The concordance between AFM and TFM readouts indicates that the SMC-like nanomechanical phenotype of LUSC-myCAFs includes both shared biological processes probed across techniques (i.e. higher actomyosin contractility) and technique-specific features (i.e. enhanced cortical stiffness in AFM or force transmission through focal adhesions in TFM). Despite this technical concordance, histotype-dependent differences were higher by AFM (70%) than by TFM (40%), which is consistent with previous reports of higher local fibroblast stiffness in response to compressive than tensile loading ^[31]^.

The hypercontractile phenotype of LUSC-myCAFs was unexpected, considering the similar α-SMA expression in LUAD and LUSC, and that pulmonary myofibroblasts are already highly contractile cells, displaying stiffness and traction forces threefold higher than TGF-β1-unstimulated fibroblasts. Mechanistically, although canonical myofibroblasts have long been described as SMC-like cells, their contractility has traditionally been attributed exclusively to NMMII motors mainly activated through the TGF-β1-RhoA/ROCK-MLCP pathway, as in non-muscle cells ^[14,15,28,34,55]^, rather than to MLCK-dependent activation of SMMII motors, which predominates in SMCs ^[41,44–46]^. In LUAD-myCAFs, their higher MYPT1^pT853^ levels ‒a canonical downstream ROCK target that suppresses MLCP activity ^[28,43]^‒, together with their nanomechanical similarity to control pulmonary myofibroblasts are consistent with a canonical non-muscle contractility program, in agreement with observations in breast and colorectal CAFs ^[34,55,57]^. In contrast, several lines of evidence support a prominent SMC-like rather than a canonical myofibroblast-like contractile program in LUSC-myCAFs, including the involvement of SMMII. First, LUSC-myCAFs showed higher expression of SM-MHC, the core component of SMMII, together with its greater colocalization with the α-SMA cytoskeleton and reduced ROCK-MYPT1 signaling. Second, LUSC-myCAFs displayed enhanced ERK1/2^pT202/Y204^, a major regulator of SMC contractility ^[47,48]^ and upstream activator of MLCK ^[29]^, extending previous observations of increased ERK1/2^pT202/Y204^ in LUSC-CAFS relative to LUAD-CAFs in the absence of TGF-β1 stimulation ^[10]^.

Intriguingly, ERK1/2 also regulates caldesmon, a classical SMC marker enriched in the LUSC stroma that was recently shown to increase cellular traction forces ^[58]^, suggesting that an ERK1/2-caldesmon axis may contribute to the nanomechanical phenotype of LUSC-myCAFs. Third, RLC^pS19/S20^ was higher in LUSC than LUAD, in line with their higher ERK1/2 activity, although the increase in phosphorylated RLC was modest compared with that of ERK1/2 and MYPT1. Since both NMMII and SMMII are activated by RLC phosphorylation and exert contractile forces through interactions with the α-SMA cytoskeleton ^[22,42,46]^, the modest increase in RLC^pS19/S20^ and the comparable α-SMA levels between histotypes are unlikely to fully account for the markedly greater contractility of LUSC-myCAFs. Yet, given the superior force-producing capacity of SMMII relative to NMMII ^[39]^, even a small increase in RLC phosphorylation could substantially enhance contractility if a fraction of the phosphorylated RLC is associated with SMMII. However, currently available RLC^pS19/S20^ antibodies do not discriminate between RLC associated with NMMII and SMMII, precluding direct assessment of the relative contribution of each motor.

TGF-β1 is a master inducer of the contractile phenotype of both canonical myofibroblasts and SMCs ^[15,51]^, and its ability to increase fibroblast stiffness has been extensively documented in normal fibroblasts ^[23,35,54]^. In pulmonary fibroblasts, previous AFM and TFM studies reported three-fold and 1.7-fold increases in stiffness and traction forces, respectively, upon TGF-β1 stimulation ^[59,60]^, consistent with our observations. In contrast, studies examining TGF-β1 effects on CAF nanomechanics are scarce and have reported either a similar stiffening to CFs in prostate cancer ^[35]^ or an attenuated stiffening in pancreatic CAFs ^[54]^. These CAF studies used either AFM or TFM alone, and provided limited mechanistic insights. We tackled these limitations in lung myCAFs by combining AFM, TFM and analyses of major contractility pathways and markers. This integrated approach revealed that TGF-β1-induced a coordinated cytoskeletal remodeling (including an increase in *MYH11* expression and stress fiber density rich in α-SMA) and enhanced actomyosin force generation, and that this contractile program was more homogeneous and amplified in LUSC-myCAFs, including increased SM-MHC expression and its colocalization with α-SMA, thereby resembling key aspects of contractile SMCs ^[41,47,50]^. Our findings also challenge the widespread use of α-SMA as a surrogate marker of TGF-β1-induced-myofibroblast contractility in culture and *in vivo* ^[15,22,23,35]^, since LUSC-myCAFs generated substantially higher contractile forces than LUAD-myCAFs despite comparable α-SMA expression and even lower *ACTA2* levels. Instead, our results highlight the importance of integrating functional nanomechanical measurements with molecular profiling, as SMC-like hypercontractile myCAFs cannot be identified by either α-SMA expression alone or current scRNAseq-based CAF classifications ^[11,24]^. In agreement with our observations, dissociations between α-SMA expression and force generation have been reported in vascular SMCs ^[61]^ and cardiac fibroblasts ^[23]^, collectively indicating that α-SMA alone is insufficient to identify SMC-like hypercontractile fibroblasts. Alternatively, the combined assessment of α-SMA and SM-MHC, including their colocalization, may provide a more accurate signature of myCAFs resembling contractile SMCs.

The causes of the non-canonical hypercontractile phenotype of LUSC-myCAFs remain unclear. Yet, the larger epigenetic alterations observed in LUSC-CAFs, evident genome-wide as global DNA hypomethylation and specifically in regulatory regions of the SMC-contractility genes *CALD1* and *MYLK*, raise the possibility that such epigenetic reprogramming reflects a stronger commitment to an SMC-like contractile phenotype. Consistently, AFM, TFM and α-SMA/SM-MHC colocalization data independently revealed reduced heterogeneity in LUSC-myCAFs, indicative of a more uniform hypercontractile phenotype. Moreover, we observed higher contractility and *MYH11* RNA levels in LUSC-CAFs than LUAD-CAFs in basal conditions (i.e. without exogenous TGF-β1), extending previous observations of higher ERK1/2^pT202/Y204^ ^[10]^, collectively revealing that LUSC-CAFs are intrinsically hypercontractile. Interestingly, larger global DNA hypomethylation has been also reported in lung CAFs from smokers compared with non-smokers ^[62]^, and pulmonary fibroblasts from smokers exhibit elevated ERK1/2 activity ^[63]^. Given the stronger association of LUSC with smoking compared to LUAD ^[5]^, which could be favored by the frequent anatomic location of LUSC within proximal airways ^[1]^, it is conceivable that both epigenetic reprogramming and smoking are involved in the selective enrichment of an SMC-like contractile phenotype in LUSC-myCAFs. However, while epigenetic regulation of both persistent myofibroblast activation and SMC plasticity are well established in organ fibrosis ^[15]^ and cardiovascular diseases ^[47]^, respectively, it remains largely unexplored in tumors ^[64]^. Likewise, future investigations should examine how the divergent DNA methylation patterns observed in upstream regulatory regions of *CALD1* and *MYLK* influence myCAF nanomechanics.

The prominent abundance of α-SMA⁺/SM-MHC⁺ myCAFs in NSCLC reported here and elsewhere ^[11]^ raises questions regarding their origin. Such cells were first described in breast cancer ^[65]^, and have also been reported in pancreatic tumors ^[11]^. Consistently, *ACTA2*⁺ *MYH11*⁺ cells have been detected by scRNAseq in NSCLC ^[57]^ and several other solid tumors ^[24]^. However, this cell population has often been interpreted as SMC rather than CAFs, based on the traditional view of *MYH11* as an exclusive SMC marker ^[15]^. In tissue repair and fibrosis, myofibroblasts are considered to arise predominantly from local mesenchymal cells, including fibroblasts, pericytes, and SMCs ^[14,15,50]^. In tumors, resident fibroblasts are regarded as the main source of myCAFs ^[21,25]^, although an earlier study reported a substantial contribution from vascular SMCs but not pericytes in breast cancer ^[65]^. In contrast, a recent pan-cancer scRNA-seq study proposed fibroblasts and pericytes, rather than SMCs, as predominant myCAF precursors across tumors, including NSCLC ^[24]^. However, the SMC-like contractile program of LUSC-myCAFs, together with the preferential location of LUSC within the proximal airways ^[1]^, ‒which are rich in vascular and airway SMCs ^[50]^‒, the well-documented plasticity of SMCs ^[50,51]^, and the evidence for SMC-derived myCAFs in breast cancer, raise the possibility that SMCs are important contributors to the myCAF population in LUSC. In contrast, LUAD-myCAFs may arise from resident fibroblasts and/or pericytes as predicted by scRNAseq analyses ^[24]^, as supported by the frequent occurrence of LUAD in the distal lung parenchyma ^[1]^, a region rich in microvasculature largely covered by pericytes ^[66]^. Supporting this interpretation, canonical pericytes generate relatively modest contractile forces ^[67]^, and rely primarily on NMMII/ROCK rather than SMMII-mediated force generation ^[68]^. Future lineage-tracing studies may clarify the contribution of true SMC-derived CAFs versus fibroblasts acquiring an SMC-like phenotype in NSCLC.

Our results add myCAF nanomechanics to the anatomic, molecular and clinical differences between LUAD and LUSC ^[1,4,5]^, providing a biomechanical framework for their divergent stromal patterns of CAF density and hypoxia. Thus, the increased SM-MHC(*MYH11*) expression and colocalization with α-SMA in LUSC-myCAFs may shift actomyosin activity toward sustained force generation at the expense of NMMII-associated dynamic migratory forces ^[26,42]^, consistent with the reduced CAF migration previously reported in LUSC ^[17]^. Contractility also correlated inversely with cell size and proliferative competence, suggesting that excessive tension impairs the growth and volume expansion required for cell-cycle progression ^[52]^. Caldesmon could reinforce this process through its ability to increase traction forces ^[58]^ and block cytokinesis ^[69]^. Interestingly, reduced proliferation and migration are also hallmarks of contractile vascular SMCs ^[50]^, reinforcing their similarity with LUSC-myCAFs, and providing a plausible biomechanical explanation for the lower CAF density observed in LUSC ^[10,17]^. Elevated mechanical forces may also compress intratumoral blood vessels and exacerbate hypoxia ^[18,19]^, consistent with our observed smaller vessel lumens and previous reports of greater hypoxia in LUSC relative to LUAD ^[4]^. Although hypoxia is often linked to tumor aggressiveness ^[6]^, it remains unclear whether our observed link between hypercontractile myCAFs and hypoxia contributes to the selective association between SMC markers and poor survival in LUSC. In addition, the emerging abundance of α-SMA⁺/SM-MHC⁺ myCAFs may have therapeutic implications, since it supports evaluating therapies that target SMC-like contractility, including antihypertensive agents ^[45]^ as well as upstream regulators such as GPCR signaling, MLCK, and ERK1/2 ^[15,45]^, particularly in LUSC. In line with this strategy, epidemiological studies have reported improved outcomes in patients receiving antihypertensive agents such as losartan across multiple solid tumors ^[18,70]^, including NSCLC ^[71]^, although histotype-specific effects have not been investigated.

## 4. Conclusions

Our study combines for the first time nanomechanical phenotyping (AFM and TFM) with contractility signaling and single-cell imaging to identify a previously unrecognized SMC-like hypercontractile program in LUSC-myCAFs. This program was characterized by enhanced ERK1/2-RLC signaling together with SM-MHC overexpression and recruitment to the α-SMA cytoskeleton, rather than by ROCK-MYPT1 signaling as in LUAD-myCAFs and canonical myofibroblasts. Our findings establish myCAF nanomechanics as a novel distinguishing feature between LUAD and LUSC, and reveal that standard SMC markers are selectively associated with adverse outcome in LUSC. Our observations support also an important role of the hypercontractile phenotype of LUSC-myCAFs in disease progression, and link it to histotype-specific patterns of stromal remodeling, including enhanced hypoxia. These findings also expose the limitations of α-SMA as a universal surrogate marker of fibroblast contractility, and demonstrate that functional nanomechanical phenotyping can uncover biologically and therapeutically relevant myCAF programs that remain hidden from conventional molecular analyses.

## 5. Experimental Section

### 5.1. Patient-derived tissue samples and fibroblasts

TMAs built from surgical NSCLC specimens (105 LUAD, 92 LUSC) were originally compiled by the CIBERES/Spanish Bronchogenic Carcinoma Cooperative Group (CIBERES cohort) ^[8]^ under approval from the Fundació Parc Taulí Ethics Committee. All TMA patients were of Caucasian ethnicity and had not received neoadjuvant therapy prior to surgery. Detailed clinicopathologic characteristics of this cohort are provided elsewhere ^[8]^ and summarized in Table S2. Primary fibroblasts were previously derived from tissue explants from a cohort of 25 NSCLC surgical patients (10 LUAD, 15 LUSC) ^[10]^, following protocols approved by the Ethics Committees of the Hospital Clínic de Barcelona. Fibroblasts were isolated in parallel from tumor tissue and matched uninvolved pulmonary tissue, the latter serving as control fibroblasts (CFs). Their mesenchymal identity was verified through vimentin positivity combined with the absence of cytokeratin staining ^[36]^. Selected patients were male, Caucasian, treatment-naïve, aged 55 or older, and current smokers; further clinical details in Table S1. Written informed consent was obtained from all patients in all cohorts, and all study procedures were in accordance with the Declaration of Helsinki.

### 5.2. Fibroblast culture and immortalization

Fibroblasts were maintained in high-glucose DMEM culture medium supplemented with 10% FBS and antibiotics as described ^[17]^. Fibroblasts from a small subset of patients were immortalized via hTERT transduction ^[7]^ (CF #5, CAF #13, CAF #20), a modification shown not to alter TGF-β1 responsiveness ^[7]^, and partially used in experiments requiring prolonged culture and/or large cell numbers (AFM, Western blotting, immunofluorescence) (Table S1). Mycoplasma-negative status was verified for all lines as reported ^[7]^. Unless stated otherwise, fibroblasts were plated on dishes pre-coated with 0.1 mg/ml type I-collagen (Ref 08-115, Sigma-Aldrich) at high density (∼9,000 cells/cm^2^), using serum-free (0% FBS) DMEM supplemented with 1% insulin-transferrin-selenium (ITS) and antibiotics (henceforth fibroblast culture medium) ^[10]^, and exposed for 3-5 days to 2.5 ng/mL recombinant human TGF-β1 (Ref 7754-BH-025, R&D Systems) for a robust myofibroblast induction or left untreated (our controls) ^[7,22]^. In these conditions, cells attain a number density within the range reported in CAFs in histologic sections ^[10,17]^ (Figure S1). This dose approximates average TGF-β1 levels reported in bronchoalveolar lavage fluid from lung cancer patients ^[16]^. For AFM experiments, cells were plated in 10% FBS fibroblast culture medium on glass-bottom dishes (35 mm, µ-Dish, Ref 80136, Ibidi). Following a 4h attachment period, medium was switched to 0% FBS fibroblast culture medium, with or without TGF-β1 for 3 days. For TFM experiments, cells were seeded in fibroblast culture medium on collagen-I-coated polyacrylamide 6 kPa gels embedded with fluorescent nanobeads, prepared as described in the TFM section. Cells were maintained in 0% FBS fibroblast culture medium with or without TGF-β1 for 5 days ^[36]^. For flow cytometry, fibroblasts were plated in fibroblast culture medium for 24h, then switched to either 0% FBS or 10% FBS fibroblast culture medium for 5 days, with a media change on day 3. For immunofluorescence, fibroblasts were seeded on collagen-coated glass-bottom 24-well plates (µ-Plate, Ref AI-82426, Ibidi) in fibroblast culture medium for 4h, and stimulated with TGF-β1 for 5 days, with a media change at day 3. Fibroblasts were cultured in fibroblast culture medium on collagen-coated 100 mm culture dishes for Western blotting or 6 well plates for qRT-PCR (Corning), with or without TGF-β1 for 20 min or 5 days, respectively. Due to the limited lifespan and availability of primary CAFs and CFs cultures, and because fibroblasts were analyzed over an extended period of time using multiple techniques, experiments were performed on the subset of cell models available at the time of analysis, resulting in partially overlapping cohorts. The specific patient cohorts used for each assay are indicated in the corresponding figure legends and summarized in Table S1.

### 5.3. TFM

Polyacrylamide gels of ∼6 kPa were prepared as described ^[36]^. Briefly, glass-bottom dishes were silanized (1:1:14 acetic acid : 3-(trimethoxysilyl)propyl methacrylate: ethanol) (Sigma), rinsed three times in ethanol, and air-dried. A gel mix of 7.5% acrylamide, 0.06% bisacrylamide, 0.5% ammonium persulfate, 0.05% TEMED (Sigma), and 1.4% 200 nm FITC-carboxylate beads (Fluospheres, Invitrogen) in 10 mM HEPES was pipetted onto the dish center (12 µl) and sandwiched under a 12 mm coverslip to polymerize. Gels were functionalized with Sulfo-SANPAH under UV light (10 min), washed with HEPES and PBS, and coated overnight at 4°C with 100 µg/ml collagen-I (Millipore). After a final PBS wash, trypsinized cells were seeded onto the gels. For TFM measurements, gel-seeded cells were imaged on an inverted microscope (Nikon Eclipse Ti) using a 20× objective, capturing phase-contrast images of cells alongside fluorescence images of the embedded beads. Cells were then trypsinized and the bead positions re-imaged in the gel’s relaxed state. Bead displacement fields (with vs. without cells) were used to generate deformation maps via custom particle-imaging-velocimetry software, from which cell traction force maps were derived using a Fourier-transform-based algorithm ^[72]^. Average traction force per unit area was calculated per cell and averaged across 9-10 cells per condition.

### 5.4. AFM

AFM experiments were performed using a JPK Nanowizard 4 BioAFM on an inverted Nikon Eclipse microscope, with triangular cantilevers bearing 5 µm wide borosilicate colloidal spheres (nominal k = 0.1 N/m; Bruker/Novascan). Optical lever sensitivity calibration was performed by acquiring force curves on a bare glass slide ^[73]^. Cantilever spring constants were calibrated via the thermal noise method in fibroblast culture medium. Nanomechanical and topographical maps (100×100 µm²) were acquired at 9 locations per sample (**Figure S**3) using JPK quantitative imaging (QI) mode (64×64 pixels, 5 µm z-length, 6 nN max force, 125 µm/s probe speed, 100 ms pixel time; <7 min/map; **Figure** S3). Raw topography images are shown unprocessed; images of the first derivative of the topography were generated using JPK’s edge-detection algorithm, and used to count stress fibers along the long cell axis ^[74]^. On average, 40 to 100 fibroblasts were measured per condition. Local Young’s modulus (*E*) was computed from the approach curves using JPK batch processing. After baseline offset and tilt correction, force versus z-piezo curves were converted to force–indentation curves, contact points were assessed automatically, and the Hertz-Sneddon model (Poisson ratio 0.5) was fitted across the full indentation range. To exclude potential mechanical artifacts due to the underlying stiff substratum, *E* data were plotted against topographic height (*h*) per condition. Some samples (8 out of 16) showed anomalously high *E* values (≥ 25 kPa) ) at very low *h* (<0.5 µm), and were excluded as outliers as reported ^[73]^ (1.9% discarded *E* values in average). No further filtering was applied. Data for figures showing force curves, profiles, and histograms were obtained through batch analysis using custom MATLAB scripts (r2024b).

### 5.5. Scanning electron microscopy

AFM cantilevers were characterized by scanning electron microscopy (SEM) prior to AFM experiments to verify microsphere geometry and surface uniformity. Cantilevers were mounted on conductive carbon tape and imaged under high vacuum using a NovaNanoSEM 230 microscope (FEI, Thermo Fisher Scientific) equipped with a through-the-lens detector.

### 5.6. Immunofluorescence

To examine the co-expression of α-SMA and SM-MHC, and corresponding morphology in single-cells, fibroblasts were fixed with 4% PFA for 15 min, permeabilized with 0.1% Triton X-100 for 10 min, stained with HCS CellMask Deep Red (Ref H32721, Thermo Fisher Scientific), and blocked for 1 h in PBS containing 10% Goat serum and 3% BSA. Primary antibodies against α-SMA (Ref M085129, Dako) and SM-MHC (MYH11) (Ref ab133567, Abcam) were co-incubated overnight at 4°C at dilutions detailed in Supporting Information. Fluorescent secondary antibodies Alexa Fluor Plus 488 goat anti-rabbit IgG (Ref A32731) and Alexa Fluor 555 goat anti-mouse IgG2a (Ref A21137) were incubated for 1h at room temperature. Nuclei were counterstained with Hoechst 33342 (Ref H3570, Invitrogen). Images were acquired using a Leica THUNDER fluorescence microscope with a 10× objective. A single-cell segmentation pipeline was developed to assess marker positivity and cell morphology, using a custom Python script (version 3.13.9; implemented using NumPy, SciPy, scikit-image, pandas, and tifffile) ^[75]^. For each image set, composite RGB images were generated by combining the Hoechst 33342 and CellMask channels for segmentation, and the Hoechst 33342, α-SMA, and SM-MHC channels for feature extraction. Single-nucleus seeds were identified from the Hoechst 33342 channel by applying Gaussian smoothing (σ = 2.5 pixels) followed by local intensity maxima detection. A foreground mask was generated from the CellMask channel using intensity thresholding combined with morphological refinement. Nuclear seeds and the foreground mask were then combined in a marker-controlled watershed algorithm (implemented using skimage.segmentation.watershed) to segment individual cells. For each segmented cell, quantitative features (spreading area and per-channel fluorescence intensity) were extracted from the corresponding feature-extraction image. Marker positivity thresholds were defined as the 99th percentile of the fluorescence intensity distribution in the corresponding negative-control samples, and were subsequently applied uniformly across all samples to determine the percentage of cells positive for each marker. Cell spreading was assessed as the projected cell area, which was computed from the segmented cell mask as the number of pixels belonging to each individual cell. Colocalization between SM-MHC and α-SMA was quantified on a per-cell basis by computing the α-SMA-in-SM-MHC Mander’s colocalization coefficient M(α-SMA/SM-MHC) as the sum of α-SMA intensity at pixels exceeding the SM-MHC threshold divided by the total α-SMA intensity within the cell ^[76]^. To analyze the α-SMA cytoskeleton alone, a similar staining protocol was used, incubating with anti-α-SMA (clone 1A4, Sigma) followed by Cy3-conjugated goat anti-mouse IgG secondary antibody (Jackson) ^[36]^. Fluorescence images were acquired on an Eclipse TE2000 microscope (Nikon) equipped with a C9100a EM-CCD camera (Hamamatsu) using a ×20 objective (n ≥ 5 random fields of view per biological repeat). α-SMA fiber intensity per cell was quantified using CSKMorphometrics software as described ^[77]^. Briefly, the software performs initial fiber segmentation, fiber refinement, and background determination/subtraction, yielding fiber intensity values for individual cells.

### 5.7. Western blotting

Proteins were extracted with a lysis buffer (50 mM Tris pH 7.4, 150 mM NaCl, 0.1% SDS, 1% Triton X-100 (Sigma), Nonidet P-40 1% (Igepal)) supplemented with protease (Cocktail Set I, Merck; Pefabloc, Roche) and phosphatase (PhosSTOP, Roche) inhibitors, and quantified with a Pierce BCA Protein Assay Kit. Equal protein amounts were resolved on Bio-Rad 4-15% precast polyacrylamide gels (90-120V, 1.5-2h) and transferred to PVDF membranes using a Bio-Rad Trans-Blot Turbo system. Membranes were blocked and incubated overnight at 4°C with shaking, using primary antibodies against ERK1/2^pT202/Y204^ (Ref 9101), total ERK1/2 (Ref 9102), MYPT1^pT853^ (Ref 4563), total MYPT1 (Ref 2634) (Cell Signaling), RLC^pS19/S20^ (Ref 600-401-416), total RLC (Ref 600-401-938, Rockland), α-SMA (Ref M085129, Dako) and α-tubulin (Ref 2144, Cell Signaling), our loading control. To examine total and phosphorylated levels of a given protein, lysates from the same samples were loaded in duplicate on two parallel gels and transferred simultaneously; one membrane was probed for the phosphorylated protein and the other for the corresponding total protein. Bound antibodies were detected with HRP-conjugated secondaries and Immobilon Forte Western HRP Substrate (Millipore), imaged on a Bio-Rad ChemiDoc system. For proteins with close molecular weights (ERK1/2, α-SMA, and α-tubulin), membranes were stripped (stripping buffer Ref 46430, Invitrogen) and reprobed with an anti-α-tubulin antibody (used as loading control). Band densitometry was performed with ImageJ, with intensities normalized to the loading control. Phosphorylated/total ratio was calculated after normalizing each signal to its loading control. Additional details in Supporting Information.

### 5.8. qRT-PCR

RNA extraction and reverse transcription were performed as described ^[4]^. Gene expression was quantified by qRT-PCR on a StepOnePlus Real-Time PCR System (Applied Biosystems), using TaqMan Gene Expression Master Mix together with TaqMan gene-specific primer/probe sets for *ACTA2* (Hs00909449_m1), *MYH11* (Hs00975796_m1), with *POLR2A* (Hs00172187_m1) serving as endogenous housekeeping gene. Relative expression with respect to the endogenous control was computed as 2^-ΔCt^ as reported ^[7]^.

### 5.9. Fibroblast number density

Cell number density in AFM experiments was assessed as reported ^[10]^. On the day of each experiment, cell samples were imaged using an inverted Leica DMIL LED microscope equipped with a Leica MC190 HD digital camera and a 5× objective (Leica Microsystems) (n ≥ 2 images/sample). Cells were counted, and corresponding number density was determined as the average cell density per calibrated image.

### 5.10. Cell Cycle

Cells were trypsinized, fixed in ice-cold 70% ethanol (Sigma) and pelleted by centrifugation (2000 rpm, 10 min). Pellets were resuspended in PBS containing propidium iodide (0.5 mg/ml) and RNase A (0.4 mg/ml, Sigma). DNA content histograms were acquired by flow cytometry (LSRFortessa, BD Biosciences) ^[37]^, and the proportion of growth-arrested fibroblasts (G0/G1 population) was determined using FACSDiva Software v6.1.3 (BD).

### 5.11. DNA methylation

DNA methylation profiles of 12 CAFs (6 LUAD, 6 LUSC) and their patient-matched CFs were obtained from the GSE68851 dataset, and generated using the Illumina Infinium 450K DNA Methylation Array ^[36]^, which provides normalized fluorescence intensities (β-values) for ∼450,000 CpGs. Relative density distributions and median values of β-values across the 1,452 CpG sites most significantly differentially methylated between CFs and CAFs (using a cutoff of absolute Δβ ≥ 0.2) were compared among CFs, LUAD-CAFs, and LUSC-CAFs to assess global DNA methylation reprogramming. Kernel density estimates were generated using the geom_density function of the *ggplot2* package in *R*. A global gene enrichment analysis was previously applied to the statistically significant differentially methylated CpG sites between CFs (n=12) and CAFs (n=12). The resulting list of genes were used to conduct KEGG pathway enrichment analysis, which reported a gene cluster annotated to the vascular smooth muscle contraction pathway (hsa04270): *PRKCE*, *KCNMA1*, *ITPR2*, *GUCY1A2*, *JMJD7-PLA2G4B*, *ITPR1*, *ADCY1*, *CALD1*, *GNAS*, *MYLK*, *MRVI1*, *GNA12*) ^[36]^. Of the four direct actomyosin regulators within this list (*MYLK*, *CALD1*, *GNA12* and *PRKCE*), only *CALD1* and *MYLK* exhibited stromal expression patterns according to the HPA (**Figure S**7). Systematic analysis of the differential DNA methylation levels between CAFs and CFs within main promoter regions and upstream regulatory sites in *MYLK* and *CALD1* identified six CpG sites with absolute differential methylation levels (|Δβ| ≥ 0.1) between CAFs and CFs within upstream regulatory regions: cg27356296 for *CALD1*, cg19534753, cg16212219, cg00465319, cg20669834, cg20050484 for *MYLK*. Corresponding CAFs/CFs methylation log10 ratios were obtained for both LUSC and LUAD.

### 5.12. Tissue microarrays (TMAs)

TMAs from the multicentric Spanish CIBERES cohort were stained for α-SMA as described ^[8]^, with nuclei counterstained using hematoxylin. Stained sections were imaged at 7-10 randomly selected fields per sample using an upright microscope (BX43) coupled to a digital camera (DP72, Olympus) with a 10× objective. Image processing and analysis were performed blind with ImageJ. Images were color-deconvoluted and binarized to calculate the positive area fraction (%), averaged per patient. TMA images from NSCLC patients (4-6 LUAD, 4-6 LUSC) stained for standard SMC contractility or differentiation markers (encoded by *MYLK*, *CALD1*, *MYH11*, *TAGLN*, *SMTN*) or angiogenesis markers (CD34, CD31) were downloaded from the HPA database (Version 19.3; http://www.proteinatlas.org) using the HPA IHC Image Downloader ^[79]^. Necrotic areas and other artifacts were excluded. SMC marker images were analyzed for % positive area as for α-SMA. Endothelial marker images were used to assess blood vessel diameter through an automated microvessel segmentation approach ^[80]^. Briefly, vessel segmentation was performed using a custom MATLAB pipeline via region-growing in HSV color space, using hues characteristic of stained endothelial cells as seeds and bright yellow-green-cyan pixels as background stop criteria. To improve segmentation accuracy, images were processed in a tiled workflow (1000×1000 pixel tiles, 250 pixel overlap). Segmented vessel objects underwent lumen detection to determine lumen perimeter and equivalent diameter, assuming circular geometry.

### 5.13. TCGA gene expression and survival analysis

Bulk *ACTA2* mRNA levels from the TCGA datasets were obtained via the UCSC Xena platform ^[81]^ (706 LUAD, 626 LUSC). Survival analysis of canonical SMC markers (*MYH11*, *TAGLN*, *SMTN*, *CALD1*, *ACTA2*) was performed using the survival and survminer R packages (version 4.4.3) ^[4]^ on the available TCGA LUAD and LUSC cohorts (498 LUAD, 486 LUSC), accessed via https://cancergenome.nih.gov ^[7]^. Briefly, patients were stratified into high- and low-expression groups for each gene using the optimal cutpoint determined by maximally selected rank statistics (maxstat R package). A similar survival analysis was conducted to evaluate the prognostic impact of combination of *ACTA2* and *MYH11* genes by combining their expression profiles.

### 5.14. Statistical analysis

Two-group comparisons were performed with two-tailed Student *t*-test unless otherwise indicated (GraphPad Prism v8.0.1). Pairwise comparisons of DNA methylation level distributions (β-values) between CFs, LUAD-CAFs, and LUSC- CAFs were performed using Wilcoxon rank-sum tests (R software). Survival curves were compared using two-sided log-rank tests in R. Statistical significance was assumed at *p* < 0.05. All data shown as mean ± SD.

## Supporting Information

Supporting Information is available from the Wiley Online Library or from the author.

## Acknowledgements

We thank Laura López (Hospital Clínic Barcelona), Patricia Fernández, Teia Vallès, Edi Reshidi, Roser Sala-Llonch (UB), Eduard Monsó (Parc Taulí) and Cooperative Lung Cancer Group CIBERES-CIBERONC-SEPAR-Biobanco for technical support and discussions.

## Author Contributions

A.C and A.B. contributed equally to this study.

A.C., A.L., M.V., M.V-M. and J.A. supervised the project.

J.A. conceptualize the project.

A.C., A.B., N.D-V., A.L., V.B., B.C. and R.M., performed the experiments.

A.C., A.B., A.L., J.R-R., M.V., L.H., M.A., M.R-P., A.P., R.M. M.K. and N.G. conducted data analysis.

A.C. and A.B. prepared the figures.

A.C., A.B., M.V-M. and J.A. wrote the manuscript. X.T., G.G. A.C., A.L., and J.A. acquired funds.

All authors contributed to the research, editing, and approval of the manuscript.

## Data Availability Statement

DNA methylation data are available in GSE68851. All other data that support the findings of this study are available from the corresponding author upon reasonable request.

