## Supplementary Information for "Nanomechanical phenotyping uncovers a non-canonical smooth muscle-like hypercontractility program in lung squamous cell carcinoma myofibroblasts"

### SUPPLEMENTARY MATERIALS

#### AFM

AFM experiments were performed using a JPK Nanowizard 4 BioAFM coupled to an Eclipse Nikon inverted optical microscope. The AFM probes used were triangular cantilevers with attached borosilicate colloidal spheres (5 μm diameter) and nominal spring constant k = 0.1 N/m (Bruker/Novascan). A glass slide was pre-cleaned before optical lever sensitivity calibration by rinsing with milliQ water and ethanol followed by dyring with pure $N_2$. Calibration was performed before measurements by collecting force curves on the bare glass slide containing fibroblast culture medium. The spring constant was calibrated using the thermal noise method after cantilever retraction from the glass slide surface while still immersed in the DMEM medium. Mechanical and topographical maps of 100×100 $\mu m^2$ size were obtained in 9 different locations for each cell sample/condition, using the JPK quantitative imaging (QI) mode in conditions that required <7 minutes/map (64×64 pixels image size, 5 μm z-length, 6 nN maximum force, 125 μm/s force curve speed, and 100 ms pixel time) (as outlined in Figure S3). Raw topography images at the measurement setpoint were represented without further processing. Images of the derivative of the raw topography were obtained with a JPK edge detection algorithm (pixel difference). The number of stress fibers was assessed by visual counting fibers along the long cell axis within topography derivative images. In average, 40-100 fibroblasts were measured by condition. The local Young´s modulus (*E*) or stiffness of each cell were computed from the approach arm of each force curve using the JPK batch processing routine. Following calibration of photodetector versus z-piezo curves, offset and tilt correction, force versus z-piezo curves were converted into force versus tip-sample distance curves. After software-based contact point assessment, the Hertz-Sneddon model was fitted throughout the entire indentation range (Poisson ratio set at 0.5). To exclude potential mechanical artifacts due to the underlying stiff substratum, *E* data were plotted against topographic height (*h*) per condition. Some samples (8 out of 16) showed anomalously high *E* values (≥ 25 kPa) ) at very low *h* (<0.5 µm), and were excluded as outliers as reported (Tello-Lafoz et al, *Immunity* 2021) (1.9% discarded *E* values in average). Sporadic $E < 0$ were also excluded. No additional filters were applied to the stiffness maps. All topography, topography derivative and stiffness images were obtained using the JPK data processing software. Plots of single force curves, topographic profiles, height and stiffness histograms were obtained with OriginPro 2022, using batch data analysis with custom Matlab algorithms (r2024b). To allow direct comparison, the force curves shown in Figure 1d and Figure 2d were collected in regions with comparatively similar height.

**Western-blot**

List of used antibodies with their corresponding blocking buffer and working dilution.

| Target | Reference | Blocking buffer | Dilution |
|---|---|---|---|
| pErk1/2$^{T202/Y204}$ | Cell Signaling, #9101 | BSA | 1:1000 |
| Erk1/2 | Cell Signaling, #9102 | Milk | 1:1000 |
| pMYPT1$^{T853}$ | Cell Signaling, #4563 | BSA | 1:1000 |
| MYPT1 | Cell Signaling, #2634 | BSA | 1:1000 |
| pRLC$^{S19/S20}$ | Rockland, 600-401-416 | BSA | 1:500 |
| RLC | Rockland, 600-401-938 | BSA | 1:500 |
| α-SMA | Dako, M085129 | Milk/BSA | 1:1000/1:100 |
| α-tubulin | Cell Signaling, #2144 | Milk | 1:2000 |

**Immunofluorescence**

List of used antibodies with their corresponding working dilution.

| Target | Reference | Dilution |
|---|---|---|
| α-SMA | Dako, clone 1A4, Ref. M085129 | 1:300 |
| SM-MHC (MYH11) | Abcam, Ref. ab133567 | 1:150 |
| Alexa Fluor Plus 488 goat anti-rabbit IgG | Invitrogen, Ref. A32731 | 1:200 |
| Fluor 555 goat anti-mouse IgG2a | Invitrogen, Ref. A21137 | 1:200 |
| Hoechst 33342 | Invitrogen, Ref. H3570 | 1:5000 |

Single-cell morphology was analyzed using quantitative morphological features extracted with skimage.measure.regionprops_table. Projected cell area was obtained from the segmented cell mask as the number of pixels belonging to each individual cell and was used as a measure of cell spreading. Additional shape descriptors included ellipse roundness and elongation ratio.

### Tissue microarrays (TMAs)

List of selected antibodies from the Human Protein Atlas.

| Protein | Selected antibody |
|---|---|
| α-SMA | CAB000002 |
| MYLK | CAB020789 |
| Caldesmon | HPA008066 |
| Transgelin | CAB001447 |
| CD34 | HPA036722 |
| CD31 | HPA004690 |
| GNA12 | CAB010046 |
| PRKCε | HPA044496 |
| SM-MHC | HPA015310 |
| Smoothelin | CAB001721 |

### SUPPLEMENTARY ACKNOWLEDGEMENTS

AC gratefully acknowledges support of the academic crowdfunding platform Academist (https://academist-cf.com/projects/251?lang=en) for the project "A touch-based microscope to detect cancer", including support from Sysmex Diagnostics S.L. together with individual supporters (Ankit Jain, Gisella Buonanno, Isidoro Calò, Andrea Calò and family, Paolo Arpino and family, Francesco Paolo Formisano, Gemma Formisano and Ortopedia Formisano srl, Gabriela Méndez de Vigo García, Simona Terminiello, Vanesa Rodríguez Osuna, Maria Chiara Biagi, Memoli Vincenzo and family, Tatsunori Ikemoto, Ryosuke Shibato, Mohit Kumar, Manish Kumar, Pratik Kumar, Prabhat Kumar and Renu Devi, Francesco Renzo, Cesare Federico Moscatelli, Roberta Caporaso, Ornella Di Somma, Pasquale and Cinzia Di Somma-Buonanno, Ida Buonanno, Antonietta Calò, Alessandra, Maria and Luigi Zurolo, Ciro Tatto, Francesco Pentangelo, Valentino and Leonilda Russo, Cristiano De Rosa, Valentina Massa and family, Marianna, Piero and Rosaria Paolino, Francesca Di Maio and family, Daniel Esteban-Ferrer, Rosaria Romano, Marina Bidello, Aurora Riccardi, Pupa Matarazzo, Amalia Dema, Alba Irollo, Valentina De Martino, Andreina Visconti, Marilisa Di Guida, Connie Pirano).

| Patient ref. | Age (y.o.) | Histologic subtype | pT | pN | Stage | AFM | TFM | IF α-SMA + MYH11 | IF α-SMA | Manders coloc. | WB | PCR *ACTA2* | PCR-*MYH11* | Cell Cycle | DNA Methy. | hTERT |
|---|---|---|---|---|---|---|---|---|---|---|---|---|---|---|---|---|
| #2 | 82 | LUSC | T3 | N2 | IIIA | | | | | | Yes | | | | | |
| #3 | 72 | LUSC | T3 | N0 | IIB | | | | Yes | | | | | | | |
| #4 | 73 | LUSC | T2b | N1* | IIB | | | Yes | Yes | Yes | | | Yes | | | |
| #5 | 69 | LUSC | T2b | N2 | IIIA | Yes | | Yes | Yes | | | | | Yes | Yes | Yes |
| #6 | 65 | LUSC | T1a | N0 | IA | Yes | | Yes | Yes | Yes | | Yes | Yes | Yes | Yes | |
| #7 | 61 | LUAD | T1b | N0 | IA | Yes | | Yes | | Yes | Yes | Yes | Yes | | | |
| #9 | 75 | LUAD | T1b | N0 | IA | | | | Yes | | | | | Yes | Yes | |
| #10 | 76 | LUAD | T1a | N0 | IA | | | Yes | | Yes | | Yes | | | | |
| #12 | 70 | LUAD | T3 | N0 | IIB | | | Yes | | Yes | Yes | Yes | Yes | Yes | Yes | |
| #13 | 59 | LUAD | T2b | N2 | IIIA | Yes | | | Yes | | Yes | Yes | Yes | Yes | Yes | Yes |
| #14 | 62 | LUSC | T3 | N1 | IIIA | | | | Yes | | | | | Yes | Yes | |
| #15 | 73 | LUAD | T1a | N0 | IA | Yes | Yes | | Yes | | | Yes | | Yes | | |
| #16 | 64 | LUSC | T2a | N1* | - | Yes | | Yes | Yes | Yes | Yes | Yes | | | Yes | |
| #17 | 80 | LUSC | T2a | N1 | IIA | | | | Yes | | | | | Yes | | |
| #18 | 83 | LUSC | T2a | N0 | IB | | | | | | | | | | Yes | |
| #20 | 78 | LUSC | T1b | N0 | IA | Yes | Yes | | | | Yes | Yes | Yes | | | Yes |
| #22 | 76 | LUSC | T2a | N1 | IIA | | Yes | | | | | Yes | | | | |
| #23 | 76 | LUSC | T2a | Nx | - | | | | Yes | | | | | Yes | | |
| #25 | 62 | LUSC | T2a | N0 | IB | | | | | | | | | | Yes | |
| #26 | 63 | LUAD | T2a | N0 | IB | | Yes | | Yes | | | Yes | | Yes | Yes | |
| #27 | 71 | LUAD | T1a | N0 | IA | Yes | Yes | | Yes | | | Yes | | | Yes | |
| #28 | 80 | LUAD | T1a | N0 | IA | | | | Yes | | | | | Yes | Yes | |
| #31 | 62 | LUSC | T3 | N0 | IIB | | | | | | | | | | | |
| #33 | 72 | LUSC | T2a | N0 | IB | Yes | Yes | | | | | Yes | | | | |
| #37 | 58 | LUAD | T1b | N0 | IA | | | | | | | Yes | | | | |

N*: direct hilar ganglionar infiltration by the tumor

**Table S1.** Summary of patient's clinical data, and their distribution across techniques

| Clinical variable | n (%) |
|---|---|
| Age (y.o.) | |
| < 65 | 90 (43.3%) |
| ≥ 65 | 118 (56.7%) |
| Gender | |
| Female | 30 (14.4%) |
| Male | 178 (85.6%) |
| Race | |
| Caucasian | 208 (100%) |
| Other | 0 |
| Smoking history | |
| Never | 20 (9.6%) |
| Former | 101 (48.6%) |
| Current | 87 (41.8%) |
| Cardiovascular comorbidity | |
| No | 118 (56.7%) |
| Yes | 90 (43.3%) |
| Histologic subtype | |
| Adenocarcinoma | 105 (50.2%) |
| Squamous cell carcinoma | 92 (44.0%) |
| Large cell carcinoma | 12 (5.7%) |
| Differentiation grade | |
| Well/mod. differentiated (G1,G2) | 151 (74.8%) |
| Poorly differentiated (G3) | 51 (25.2%) |
| Tumor stage (TNM)[a] | |
| IA1 | 6 (2.9%) |
| IA2 | 21 (10.2%) |
| IA3 | 31 (15.0%) |
| IB | 47 (22.8%) |
| IIA | 15 (7.3%) |
| IIB | 56 (27.2%) |
| IIIA | 29 (14.1%) |
| IVA | 1 (0.5%) |

*NOTE:* [a]According to 8$^{th}$ edition of the IASLC staging classification;

**Table S2.** Summary of clinicopathologic variables of the lung cancer patients from the CIBERES cohort used in the TMAs

### SUPPLEMENTARY FIGURES

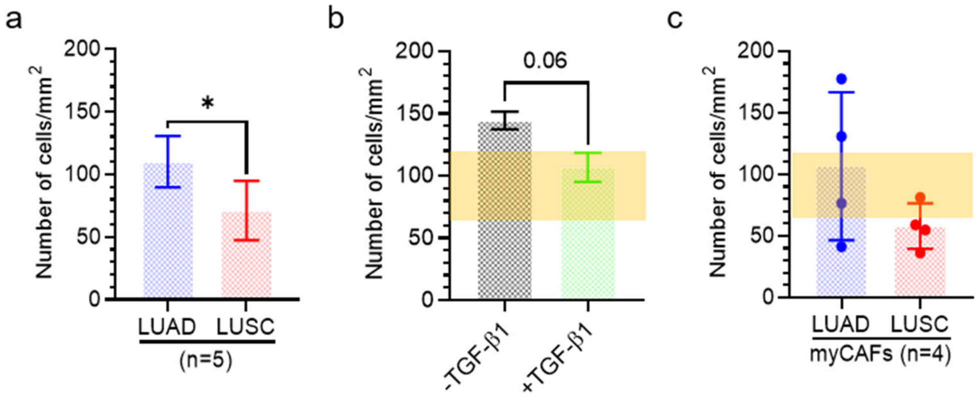


**Figure S1.**

CAF number density in LUAD and LUSC. (a) CAF number density assessed by morphometric analysis of hematoxylin-eosin images of surgical NSCLC patients (5 LUAD, 5 LUSC). (b) Fibroblast number density of CFs (#5) cultured as in Figure 1d. (c) Fibroblast number density of myCAFs cultured as in Figure 2d (4 LUAD, 4 LUSC). A semitransparent horizontal band covering the mean ± SD CAF number density (65 ± 116 cells/mm$^2$) from a) was added in (b) and (c) as a reference of the high CAF density observed in histologic staining from patient biopsies. Error bars represent mean ± SD. * $p < 0.05$; **, $p < 0.01$; ***, $p < 0.001$. Two-group statistical comparisons were done using two-tailed Student's t test.

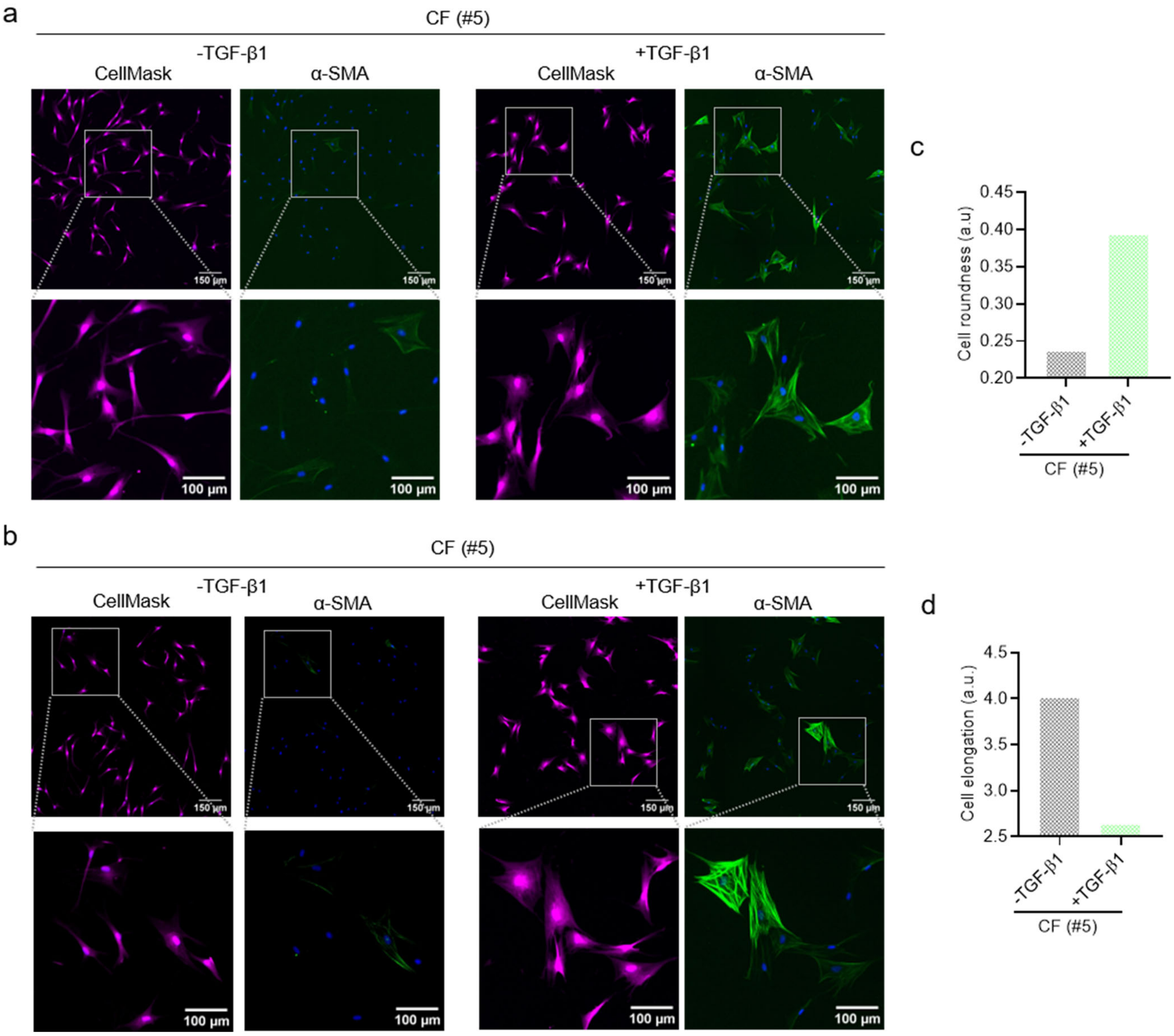


**Figure S2.**

TGF-β1 impact on α-SMA expression and cell morphology in control pulmonary fibroblasts. (a-b) Representative full-field fluorescence images of CellMask and α-SMA merged with Hoechst 33342 staining of control pulmonary fibroblasts (CFs; patient #5) cultured in the absence (-TGF-β1) or presence of TGF-β1 (+TGF-β1) for 5 days. Selected regions are shown in bottom panels. (c) Cell roundness of CFs (#5) cultured as in (a) assessed from morphometric analysis of CellMask images. (d) Cell elongation ratio of CFs (#5) cultured as in (a) assessed from morphometric analysis of CellMask images.

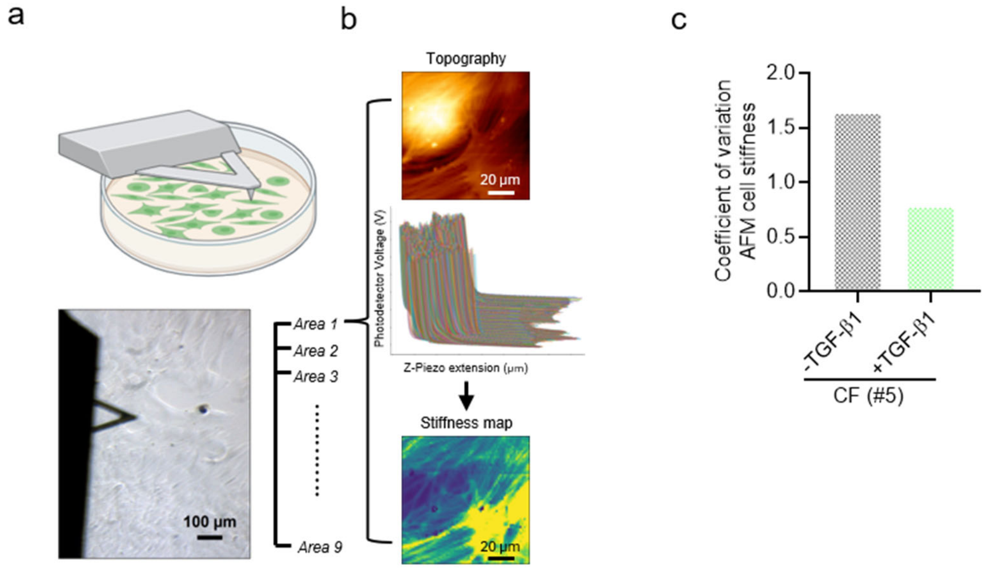


**Figure S3.**

Outline of AFM experiments and data analysis. (a) (top) Cartoon of a representative AFM probe used to measure cultured fibroblasts; (bottom) phase contrast image of an actual AFM probe on top of myCAFs. (b) Schematic of the workflow used to analyze AFM measurements obtained in 9 regions within each sample. Top panel shows a topographic image of LUSC-myCAFs (#6) collected in quantitative imaging mode. Middle panel shows raw photodetector voltage (proportional to the force (*F*)) vs. z-piezo. Fittings to the Hertz-Sneddon model yeilds the Young´s modulus (*E*) used to generate the corresponding stiffness maps (bottom panel). (c) Coefficient of variation (mean/SD) of *E* data obtained on CFs (#5) cultured in the absence or presence of TGF-β1 for 3 days.

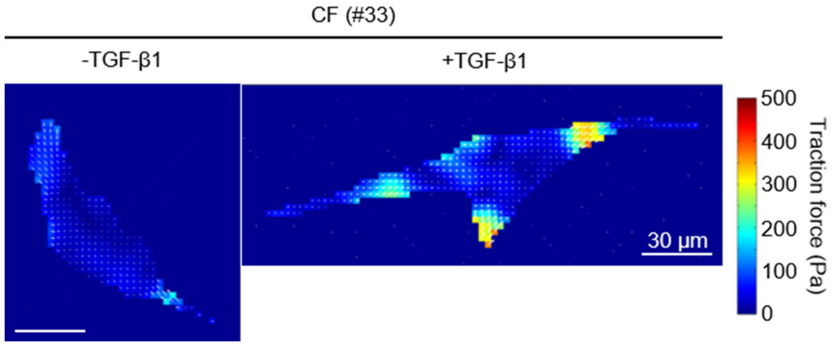


**Figure S4.**
Representative maps of traction forces obtained with TFM on a single CF (#33) cultured in the absence or presence of TGF-β1 for 5 days.

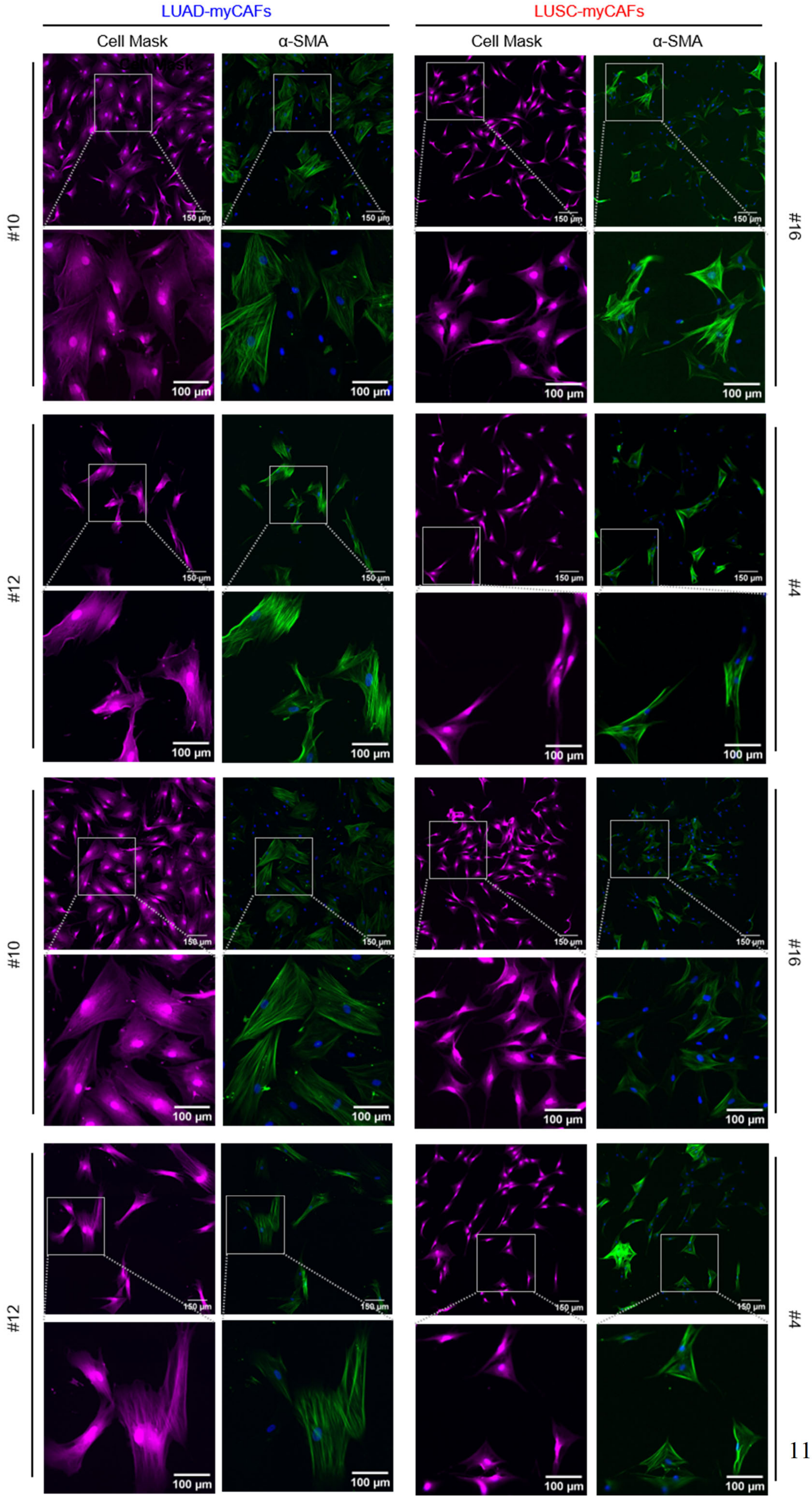
LUAD-myCAFs
LUSC-myCAFs
Cell Mask
α-SMA
Cell Mask
α-SMA
#10
#12
#10
#12
#16
#4
#16
#4
150 µm
100 µm

**Figure S5.**

Representative full-field fluorescence images of CellMask and α-SMA merged with Hoechst 33342 staining of LUAD-myCAFs (left panels; #10, #12) and LUSC-myCAFs (right panels; #4, #16) stimulated with TGF-β1 for 5 days. Selected regions are shown in corresponding bottom panels.

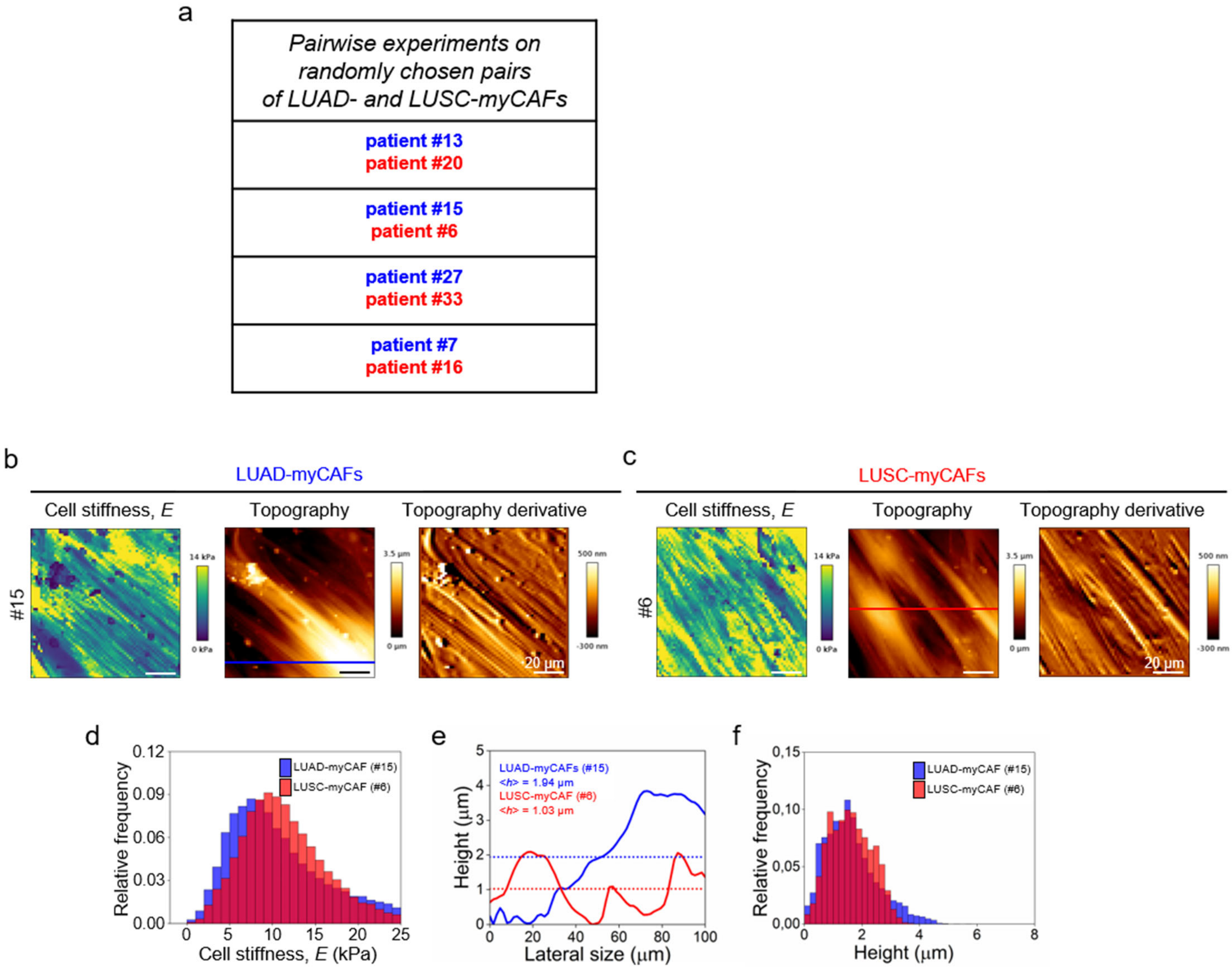


**Figure S6.**

List of histotype-pairs and representative AFM measurements on myCAFs. (a) List of LUAD-LUSC histotype pairs measured within the same experiment. (b) Representative stiffness map (left), and corresponding topography image (middle) and its first derivative (right) measured on a single region in LUAD-myCAFs (#15) cultured as in Figure 2d, which correspond to one of the 4 LUAD-LUSC histotype pairs analyzed in the same experiment (a). (c) Representative stiffness map (left), and corresponding topography image (middle) and its first derivative (right) measured on a single region in LUSC-myCAFs (#6) cultured as in (b). (d) Distributions of *E* values measured on LUAD-myCAFs (#15) and LUSC-myCAFs (#6) cultured as in (b) (9 regions per sample). *E* data were processed as in Figure 1f. (e) Height (*h*) profiles along the horizontal lines indicated in the topography images in (b-c). Dashed lines indicate the corresponding average height (<*h*>) across the profile. (f) Distributions of height values of myCAFs cultured as in (b) (9 images/sample) from the histotype pair shown in (b-c).

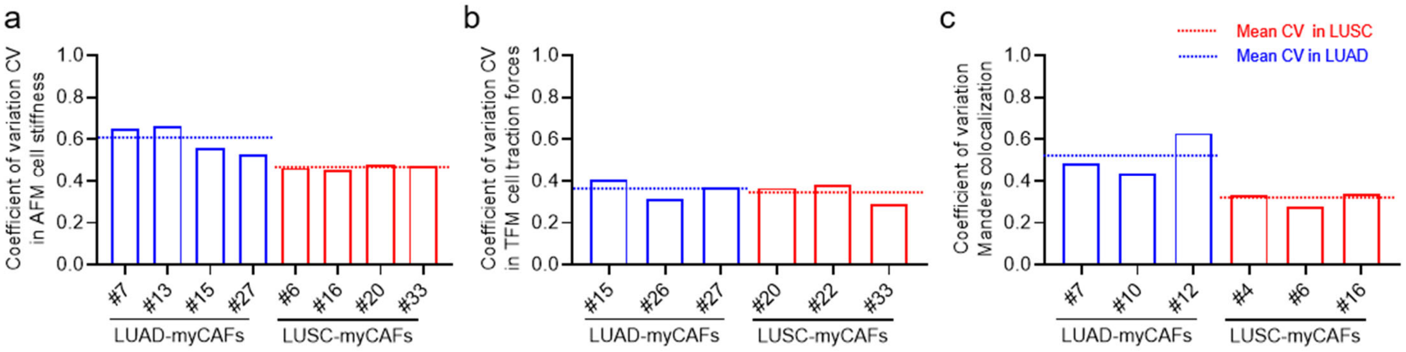


**Figure S7.**

Variability of AFM, TFM and Manders colocalization data. (a) Coefficient of variation (*CV* = mean/SD) of all *E* data obtained in myCAFs from each patient (4 LUAD, 4 LUSC) and processed as in Figure 1f. (b) *CV* of the average cell traction forces obtained in myCAFs from each patient (3 LUAD, 3 LUSC). (c) *CV* of the average cell Manders colocalization coefficient M(α-SMA/SM-MHC) obtained in myCAFs from each patient (3 LUAD, 3 LUSC). The average *CV* values of LUAD-myCAFs and LUSC-myCAFs are shown as a blue and red dashed horizontal lines, respectively (a-c).

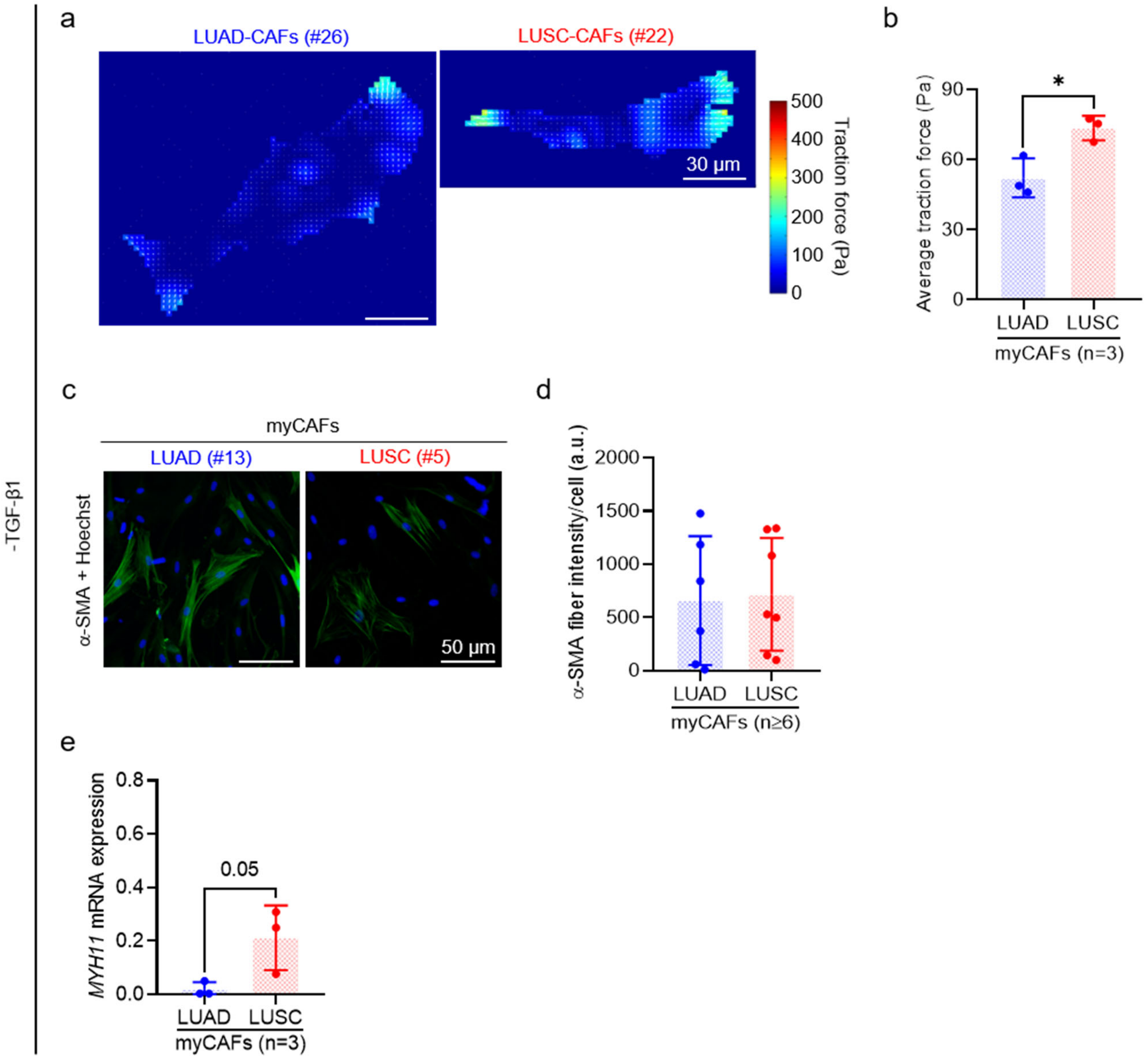


**Figure S8.**

Nanomechanical characterization of CAFs in basal conditions (absence of exogenous TGF-β1). (a) Representative maps of traction forces obtained with TFM on a single LUAD-CAF (left) and LUSC-CAF (right) cultured in the absence of TGF-β1 for 5 days. (b) Average cell traction forces of CAFs cultured as in (a). Each dot represents a single patient (3 LUAD, 3 LUSC). (c) Representative images of α-SMA merged with Hoechst 33342 nuclear fluorescence staining of CAFs cultured as in (a). (d) α-SMA fiber intensity/cell in CFs cultured as in (a). Each dot represents a single patient (6 LUAD, 7 LUSC; listed in Table S1). (e) mRNA levels of *MYH11* assessed by qRT-PCR on CAFs cultured as in (a) (3 LUAD, 3 LUSC; listed in Table S1). Error bars represent mean ± SD. * $p < 0.05$; **, $p < 0.01$; ***, $p < 0.001$. Statistical comparisons were done using two-tailed Student's t test.

a

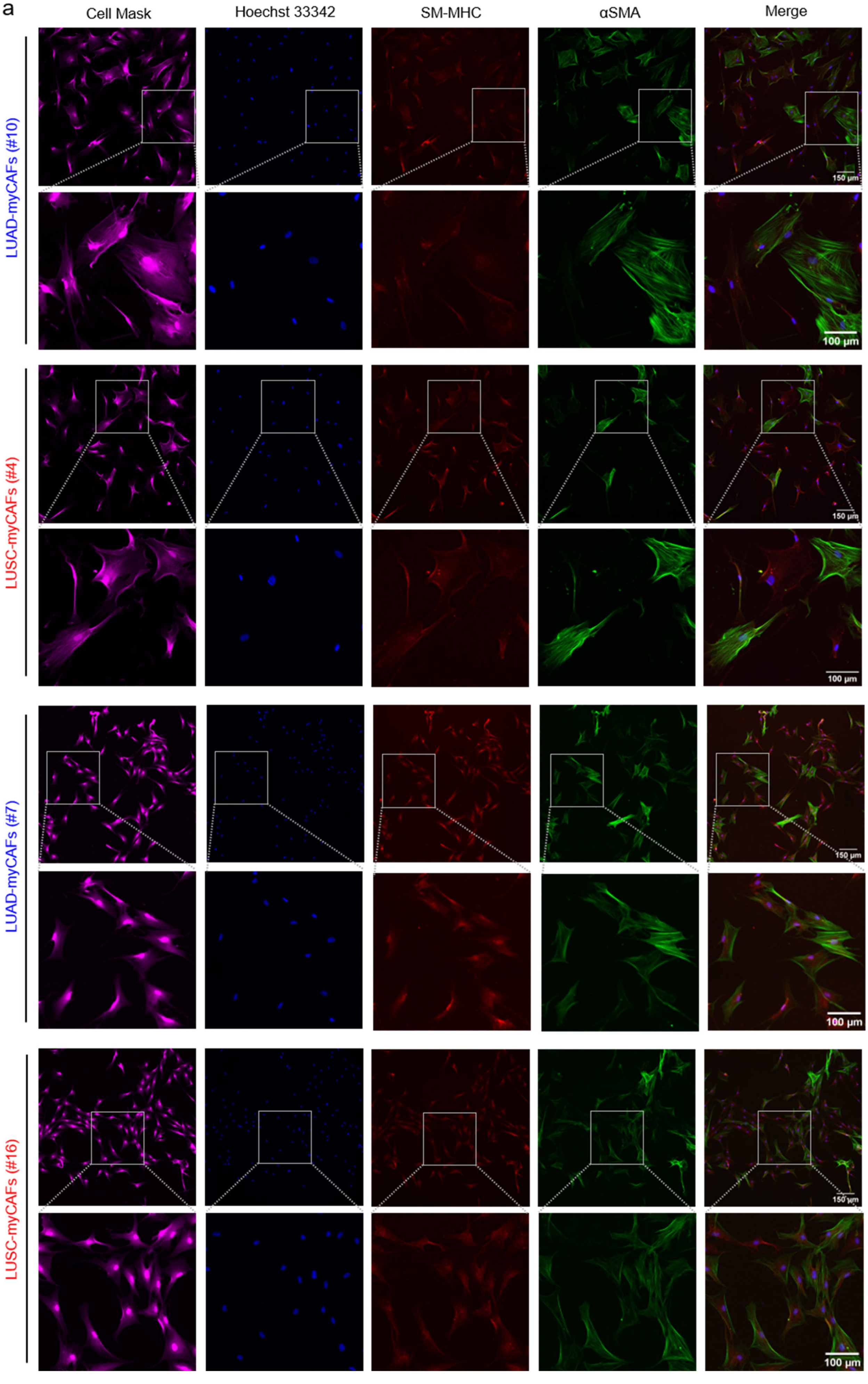

Cell Mask
Hoechst 33342
SM-MHC
αSMA
Merge
LUAD-myCAFs (#10)
LUSC-myCAFs (#4)
LUAD-myCAFs (#7)
LUSC-myCAFs (#16)
150 μm
100 μm

**Figure S9.**

Representative full-field fluorescence images of CellMask (1st column), Hoechst 33342 ($2^{nd}$ column), SM-MHC ($3^{rd}$ column), α-SMA ($4^{th}$ column) and merged SM-MHC and α-SMA channels ($5^{th}$ column) of LUAD-myCAFs (#7, #10) and LUSC-myCAFs (#4, #16) stimulated with TGF-β1 for 5 days. Selected regions are shown in corresponding bottom panels.

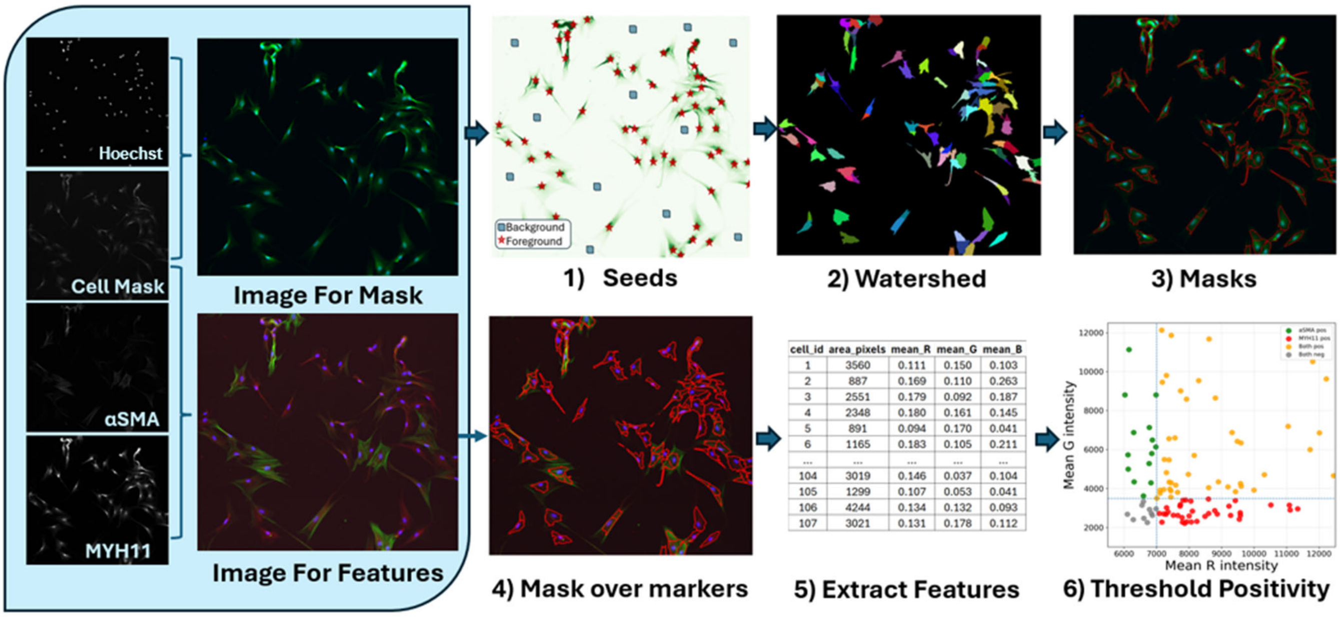


**Figure S10.**

Details of the CellMask-based single-cell fluorescence analysis of SM-MHC and α-SMA co-staining in myCAFs. A cell segmentation pipeline was developed for multi-channel fluorescence images using a Python script. First, two RGB images were generated to separate the channels used for segmentation (Hoechst 33342 and CellMask) and for feature extraction (Hoechst 33342, α-SMA, and SM-MHC (*MYH11*)). Nuclei seeds were detected from the Hoechst 33342 channel after Gaussian smoothing by identifying local intensity peaks. A foreground mask was then generated from the CellMask channel using intensity thresholding and morphological refinement. These seeds and the foreground mask were combined in a marker-controlled watershed algorithm to segment individual cells. For each segmented cell, quantitative features were extracted from the feature extraction RGB image, including area and channel intensity measurements.

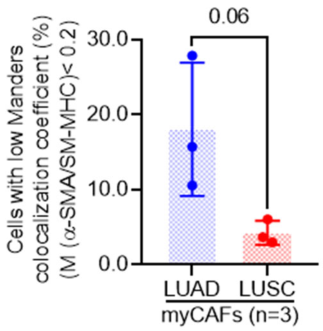


**Figure S11.**

Percentage (%) of cells with low Manders colocalization coefficient (M (α-SMA/SM-MHC) < 0.2). Each dot represents a single patient (3 LUAD, 3 LUSC; listed in Table S1). Error bars represent mean ± SD. * $p < 0.05$; **, $p < 0.01$; ***, $p < 0.001$. Statistical comparisons were done using two-tailed Student's t test.

a

| Protein (*Gene*) | Stroma score (*) | Cancer cell score (*) |
|---|---|---|
| Caldesmon (*CALD1*) | 2 | 0 |
| Guanine nucleotide-binding protein subunit alpha-12 (*GNA12*) | 0 | 2 |
| Myosin light chain kinase (*MYLK*) | 1 | 0 |
| Protein kinase C epsilon (*PRKCE*) | 0 | 0 |
| SM-MHC (*MYH11*) | 1 | 1 |
| Smoothelin (*SMTN*) | 1 | 1 |
| Transgelin (*TAGLN*) | 1 | 0 |
| α-SMA (*ACTA2*) | 2 | 0 |

(*) Histoscore: 0 negative; 1 weak; 2 strong

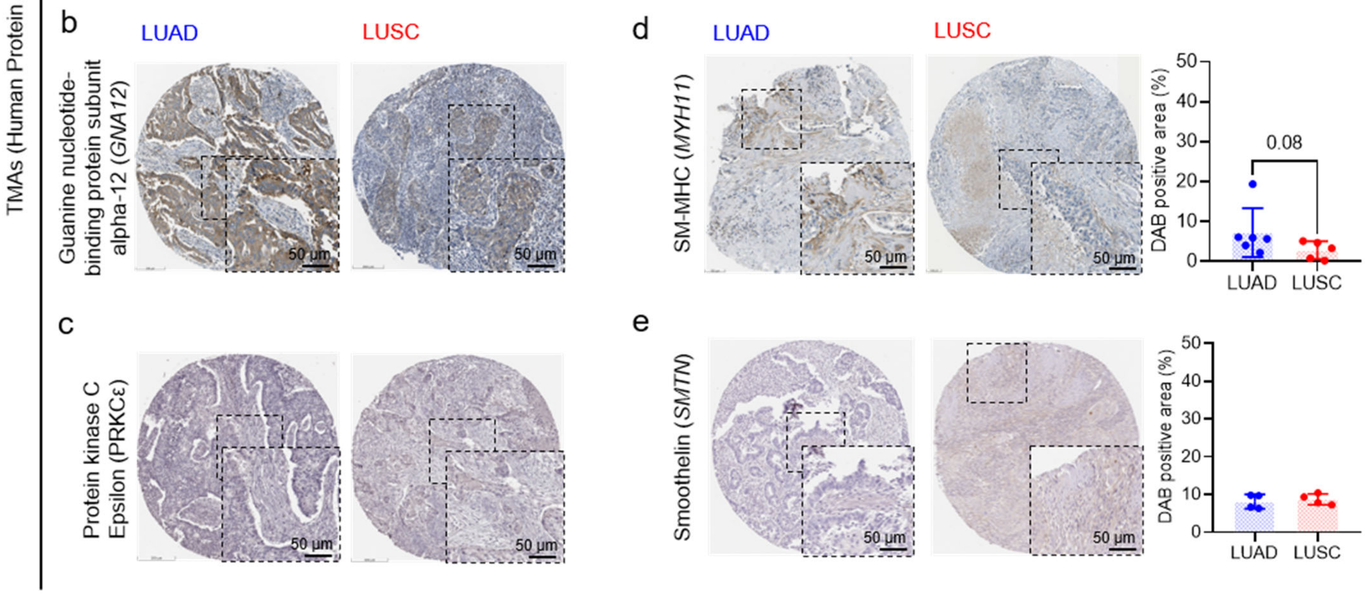


**Figure S12.**

Histologic analysis of smooth muscle cell (SMC)-associated contractility genes in TMAs from the Human Protein Atlas. (a) Histoscore of a panel of SMC-associated contractility regulatory genes (annotated to the KEGG vascular smooth muscle contraction pathway (hsa04270): *MYLK*, *CALD1*, *GNA12* and *PRKCE*) and standard SMC markers (*MYH11*, *SMTN*, *TAGLN*, *ACTA2*). Marker histologic scoring within the stromal and cancer compartment was based on three categories (negative-0, weak-1, and strong-2), under the supervision of lung cancer pathologist Dr Laura López at Hospital Clínic Barcelona. (b) Representative images of guanine nucleotide-binding protein subunit alpha-12 (encoded by *GNA12*) histological staining in TMAs from LUAD and LUSC tumors obtained from the HPA. Inset shows a magnified view of the indicated region, showing a prominent staining in the cancer rather in the stromal compartment. (c) Representative images of protein kinase C epsilon (encoded by *PRKCE*) histological staining in TMAs from LUAD and LUSC tumors obtained from the HPA. Inset shows a magnified view of the indicated region, showing largely negative staining in both the cancer and stromal compartments. (d) (left) Representative images of SM-MHC (encoded by *MYH11*) histological staining in TMAs from LUAD and LUSC tumors obtained from the HPA (antibody HPA015310). Inset shows a magnified view of the indicated region; (right) corresponding average quantification of the percentage (%) of positively stained area per patient (6 LUAD, 5 LUSC). (e) (left) Representative images of smoothelin (encoded by *SMTN*) histological staining in TMAs from LUAD and LUSC tumors obtained from the HPA (antibody

CAB001721). Inset shows a magnified view of the indicated region; (right) corresponding average quantification of the percentage (%) of positively stained area per patient (4 LUAD, 4 LUSC). Error bars represent mean ± SD. * $p < 0.05$; **, $p < 0.01$; ***, $p < 0.001$. Statistical comparisons were done using two-tailed Student's t test.

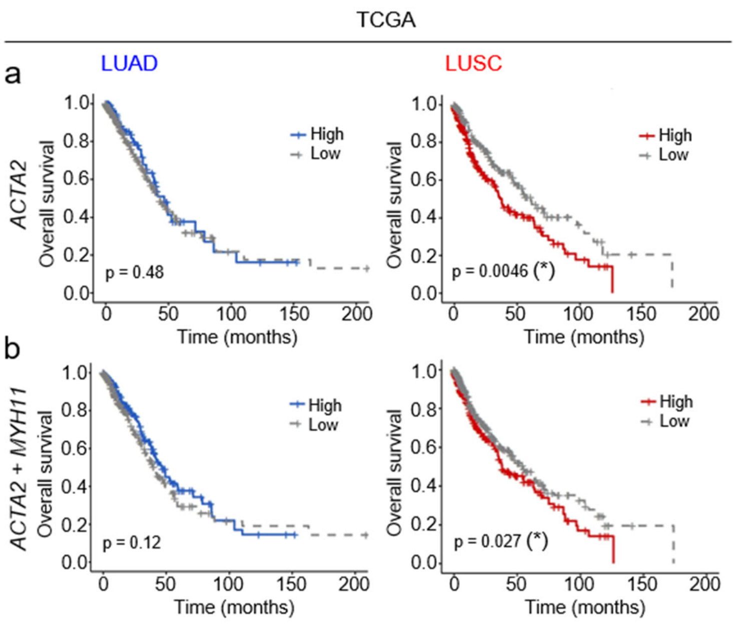


**Figure S13.**

Survival curves of contractility markers *ACTA2* and *MYH11*. (a-b) Kaplan-Meier survival curves stratifying patients according to standard SMC contractility markers, including *ACTA2* alone (a) or combined with *MYH11* (b). Survival data obtained from TCGA dataset (498 LUAD, 486 LUSC). (*) indicates significant p-value. * $p < 0.05$; **, $p < 0.01$; ***, $p < 0.001$ comparing survival curves by log-rank test.
